# Improvement of cell growth in green algae *Chlamydomonas reinhardtii* through co-cultivation with yeast *Saccharomyces cerevisiae*

**DOI:** 10.1101/2023.09.27.559874

**Authors:** Yukino Karitani, Ryosuke Yamada, Takuya Matsumoto, Hiroyasu Ogino

**Affiliations:** Osaka Metropolitan University, Department of Chemical Engineering, 1-1 Gakuen-cho, Naka-ku, Sakai, Osaka 599-8531, Japan

**Keywords:** *Chlamydomonas reinhardtii*, Co-cultivation, Photosynthesis, *Saccharomyces cerevisiae*, Transcriptome analysis

## Abstract

Biological fixation methods have attracted considerable attention because they can be applied for the fixation of dilute CO_2_ in the atmosphere. Co-cultivation of certain microalgae with heterotrophic microorganisms can increase the growth potential of microalgae under dilute CO_2_ conditions. The objective of this study was to determine the culture conditions under which the growth potential of green algae *Chlamydomonas reinhardtii* is enhanced by co-culturing with the yeast *Saccharomyces cerevisiae*, and to identify the cause of the enhanced growth potential using transcriptome analysis. When *C. reinhardtii* and *S. cerevisiae* were co-cultured with an initial green algae to yeast inoculum ratio of 1:3, the cell concentration of *C. reinhardtii* reached 133 × 10^5^ cells/mL on day 18 of culture, which was 1.5 times higher than that of the monoculture. Transcriptome analysis revealed that the expression levels of 363 green algae and 815 yeast genes were altered through co-cultivation. These include genes responsible for ammonium transport and CO_2_ enrichment mechanism in green algae and the genes responsible for glycolysis and stress responses in yeast. In conclusion, we identified the culture condition suitable for the co-cultivation of *C. reinhardtii* and *S. cerevisiae*. In addition, we discuss the cause of the increased growth potential of *C. reinhardtii* based on transcriptome analysis data. Although further studies are needed to elucidate the full impact of microbial interactions in *C. reinhardtii* and *S. cerevisiae* co-cultures, the findings of this study represent an important first step toward achieving this goal.

## 1. Introduction

Against the background of global warming, there is a need to fix and reduce CO_2_ emissions, which have a greenhouse effect [1]. CO_2_ fixation technologies mainly include physical, chemical, and biological methods. Biological fixation is a CO_2_ fixation method that utilizes photosynthesis in plants and algae [2]. Physical methods are mainly applied to high-pressure and high-concentration CO_2_; thus, they are unsuitable for fixing dilute CO_2_ in the atmosphere [3]. However, the biological fixation method attracts a lot of attention because it can be applied not only to the fixation of high-concentration CO_2_ but also to the fixation of dilute CO_2_ in the atmosphere [4].

Green algae, which are autotrophic microorganisms that perform photosynthesis, grow using CO_2_ as the sole carbon source; therefore, it is possible to fix CO_2_ by culturing green algae. In addition, because green algae accumulate excess carbon as starch, glycogen, and lipids in their cells, studies are underway to use green algae as food and feed [5]. In addition, green algal cells can be used as microbial fertilizers instead of chemical fertilizers, which have high environmental impacts [6]. Moreover, their growth rates and photosynthetic efficiencies are higher than those of terrestrial plants; therefore, biological CO_2_ fixation by culturing green algae has attracted considerable attention [7]. However, green algae generally exhibit low CO_2_ fixation efficiency under CO_2_ dilute atmospheric conditions. Therefore, efficient CO_2_ fixation by green algae requires high concentrations of CO_2_ [8].

Co-cultivation of green algae, which are autotrophic microorganisms, and heterotrophic microorganisms, which require an organic carbon source for growth, in a specific combination in an air atmosphere, makes it possible to improve the growth ability and metabolite productivity of both microorganisms. In previous studies, the co-cultivation of the green alga *Chlamydomonas reinhardtii* and *Escherichia coli*, the green alga *Scenedesmus obliquus* and the yeast *Candida tropicalis*, and the green alga *Chlorella pyrenoidosa* and the yeast *Rhodotorula glutinis* have been investigated [9-11]. In these co-culture systems, various metabolites are exchanged between the two types of microorganisms, and oxygen consumption by heterotrophic microorganisms alleviates oxygen inhibition that hinders the growth of green algae. However, the detailed mechanism by which co-culture improves the growth potential has not been clarified. The optimal combination of microorganisms in co-culture, changes in gene expression due to co-culture, and the metabolites exchanged between the two microorganisms are not fully understood. Further investigation of the interaction between the two microorganisms in co-culture is expected to establish a co-culture method that achieves a high growth potential of green algae and realizes highly efficient dilute CO_2_ fixation.

Among green algae, *C. reinhardtii* has a relatively high growth rate under atmospheric conditions compared with other common green algae [12]. In addition, the entire genome of *C. reinhardtii* has been decoded and a genetic recombination method for *C. reinhardtii* has been established [13]. Furthermore, *C. reinhardtii* is a generally recognized as safe (GRAS) microorganism, and its use in food and medicine is under consideration [14]. Moreover, the yeast *Saccharomyces cerevisiae* is also a safe GRAS microorganism, and its cultivation technology has been established on an industrial scale [15]. It is also easily genetically modified and produces a variety of useful compounds from sugars such as glucose obtained by starch degradation [16]. Therefore, if an efficient co-culture system between *C. reinhardtii* and *S. cerevisiae* could be established, it would have a wide range of applications. However, there have been few examples of co-cultivation of *C. reinhardtii* and *S. cerevisiae*.

The objective of this study was to determine the culture conditions under which the growth potential of *C. reinhardtii* is enhanced by co-culturing with *S. cerevisiae*, and to identify the cause of the enhanced growth potential. First, *C. reinhardtii* and *S. cerevisiae* were cultured at various initial inoculum ratios to evaluate the effects of the initial inoculum ratio on the growth of each microorganism. Transcriptome analysis of *C. reinhardtii* and *S. cerevisiae* was performed under culture conditions that increased the growth potential of *C. reinhardtii* and the reasons for the increased growth potential of *C. reinhardtii* are discussed.

## 2. Materials and Methods

### 2.1. Strains and media

*C. reinhardtii* NIES-2238 (National Institute for Environmental Studies) and *S. cerevisiae* YPH499 (NBRC 10505; NITE Biological Resource Center) are the strains used in the study. BG11Y medium (1 vol% BG11 broth for microbiology [Sigma-Aldrich Japan, Tokyo, Japan], 5 g/L yeast extract [Formedium, Norfolk, UK], and 0.1 vol% trace metal mix A5 + Co [Sigma-Aldrich Japan]) and yeast/peptone/dextrose (YPD) medium (10 g/L yeast extract, 20 g/L peptone [Formedium], and 20 g/L glucose [Nacalai Tesque, Kyoto, Japan]) were used for culturing. For solid media, 20 g/L of agar powder (Nacalai Tesque) was added.

### 2.2. Co-culture of green algae and yeast

Monoculture and co-culture of green algae and yeast were conducted in 50 mL of BG11Y medium in a 250 mL flask. The flasks were shaken at a constant light intensity of 60 μmol-photons/m^2^/s, 30°C, and 120 rpm using a shaker installed in a growth chamber (CLE-305; Tomy Seiko, Tokyo, Japan) under atmospheric conditions.

On one hand, the initial cell concentration of green algae was 2.5 × 10^5^ cells/mL. On the other hand, the initial cell concentration of yeast was 2.5 × 10^5^, 8.1 × 10^5^, 16 × 10^5^, 32 × 10^5^, 64 × 10^5^, 128 × 10^5^, and 255 × 10^5^ cells/mL (equivalent to 1, 3, 6, 13, 25, 51, and 101 times the cell density of green algae, respectively). Pre-cultures were performed in YPD medium (yeast) for 3 d or in BG11Y medium (green algae) for 7 d at a constant light intensity of 60 μmol-photons/m^2^/s, 30°C, and 120 rpm in a flask under atmospheric conditions. Light intensity was measured using an optical analyzer (LA-105, Nippon Medical & Chemical Instruments, Osaka, Japan).

### 2.3. Analysis of cell concentration and chlorophyll

The cell concentration was measured using a flow cytometer (CyFlow Cube 6, SYSMEX CORPORATION, Kobe, Japan) with absolute number counting function, and the number of cells contained in 200 μL of cell suspension flowed at 2.0 μL/s. Prior to the measurement, 25% (w/v) glutaraldehyde (Nacalai Tesque) was added to a final concentration of 2 vol%, shaken at 37°C and 1200 rpm for 2 h, and stained with SYTOX Green (Thermo Fisher Scientific, MA, USA).

The chlorophyll content was measured as previously described [11]. The absorbances at 665 and 650 nm were recorded using a spectrophotometer (UVmini-1240, Shimadzu, Kyoto, Japan).

Chlorophyll content was calculated using equation (1).

Chlorophyll a (mg/L) = 16.5 × ABS_665_ - 8.3 × ABS_650_ (1)

### 2.4. Transcriptome analysis

Total RNA was isolated from microbial cells using NucleoSpin RNA (Takara Bio, Otsu, Japan). To prepare a complementary DNA (cDNA) library for next generation sequencing from the extracted RNA, MGIEasy RNA Directional Library Prep Set (MGI Tech, Shenzhen, China) was used. RNA sequencing was performed using DNBSEQ-G400 (MGI Tech).

The genome sequences of *C. reinhardtii* CC-503 and *S. cerevisiae* S288c were used as reference sequences for read mapping using Geneious prime version 2020.0.3 (Tomy Digital Biology, Tokyo, Japan). Differentially expressed genes (DEGs) were identified by calculating differential expression log_2_ ratios and p-values using Geneious.

## 3. Results

### 3.1. Monoculture and co-culture of green algae and yeast

Monocultures and co-cultures of green algae and yeast in BG11Y medium were conducted. The initial cell concentration of green algae was 2.5 × 10^5^ cells/mL, and that of yeast was 1, 3, 6, 13, 25, 51, and 101 times the initial cell concentration of green algae.

The time course of green algae cell concentration is shown in Fig. 1(a). The green algal cell concentration after 18 d of culture is shown in Fig. 1(b). The concentration of green algae in the monoculture tended to increase until 18 d of cultivation, reaching 91 × 10^5^ cells/mL. Similarly, the cell concentration of green algae in the co-culture tended to increase until 18 d of cultivation and was higher than that in the monoculture under all initial cell inoculation conditions. In particular, when the ratio of green algae to yeast was 1:3, the cell concentration reached 133 × 10^5^ cells/mL, which was the highest value among all conditions after 18 d of culture. This was 1.5 times higher than that in the monoculture (91 × 10^5^ cells/mL).

**Fig. 1.**
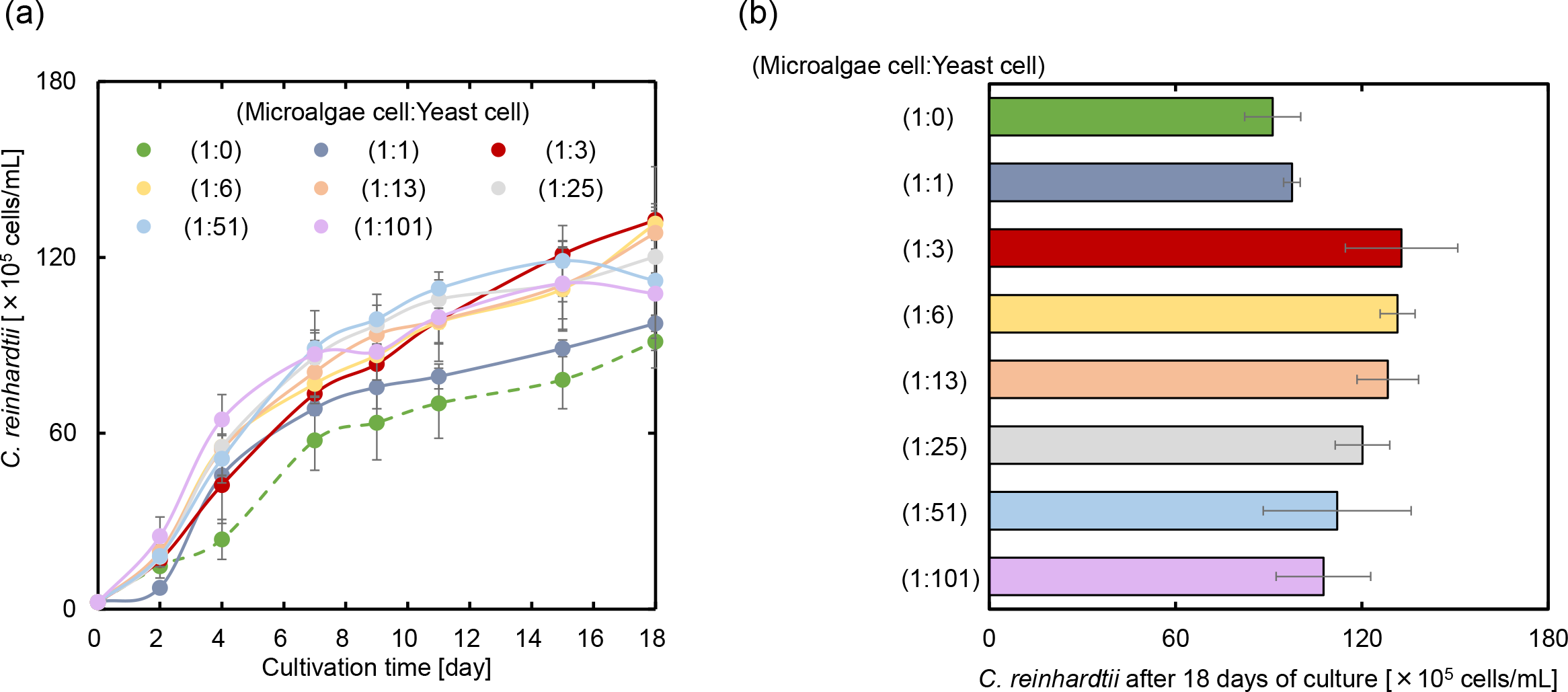
(a) Time course of green algae cell concentration and (b) green algae cell concentration after 18 d of culture. Three biological replicates were performed and all data reported as mean ± standard deviation.

The time course of yeast cell concentration when the ratio of green algae to yeast was 0:3 and 1:3 is shown in Fig. 2(a). The yeast cell concentration after 18 d of culture is shown in Fig. 2(b). Both the yeast monoculture and co-culture exhibited similar growth behavior for up to 4 d of culture. However, the yeast monoculture grew better after that, reaching cell numbers 230 × 10^5^ and 107 × 10^5^ cells/mL in monoculture and co-culture, respectively, after 18 d of culture.

**Fig. 2.**
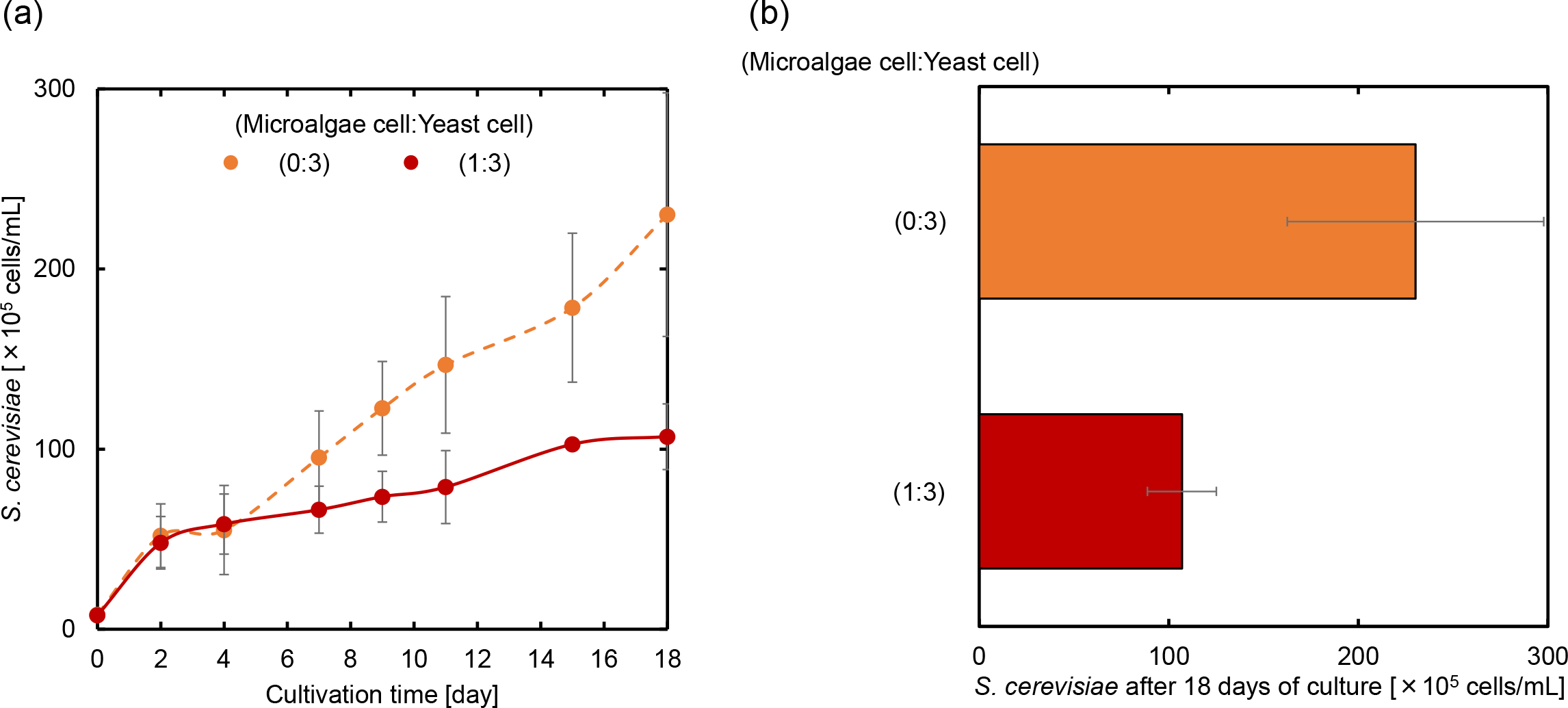
(a) Time course yeast cell concentration and (b) yeast cell concentration after 18 d of culture. Three biological replicates were performed and all data reported as mean ± standard deviation.

The time course of chlorophyll concentrations in the co-culture and green algae monoculture is shown in Fig. 3(a). The chlorophyll concentration after 18 d of culture is shown in Fig. 3(b). The chlorophyll concentration increased in the green algal monoculture until 18 d of culture, reaching 7.6 μg/mL. Similarly, the chlorophyll concentration increased in the co-culture until 18 d after cultivation and was higher than that in the monoculture under all initial cell inoculation conditions. When green algae and yeast were co-cultured, the higher the initial inoculum concentration of yeast, the higher the chlorophyll concentration was. The chlorophyll concentration reached 17.9 μg/mL after 18 d of co-cultivation at an inoculum ratio of green algal cells and yeast cells of 1:101. This value was 2.4 times higher than that of the monoculture (7.6 μg/mL).

**Fig. 3.**
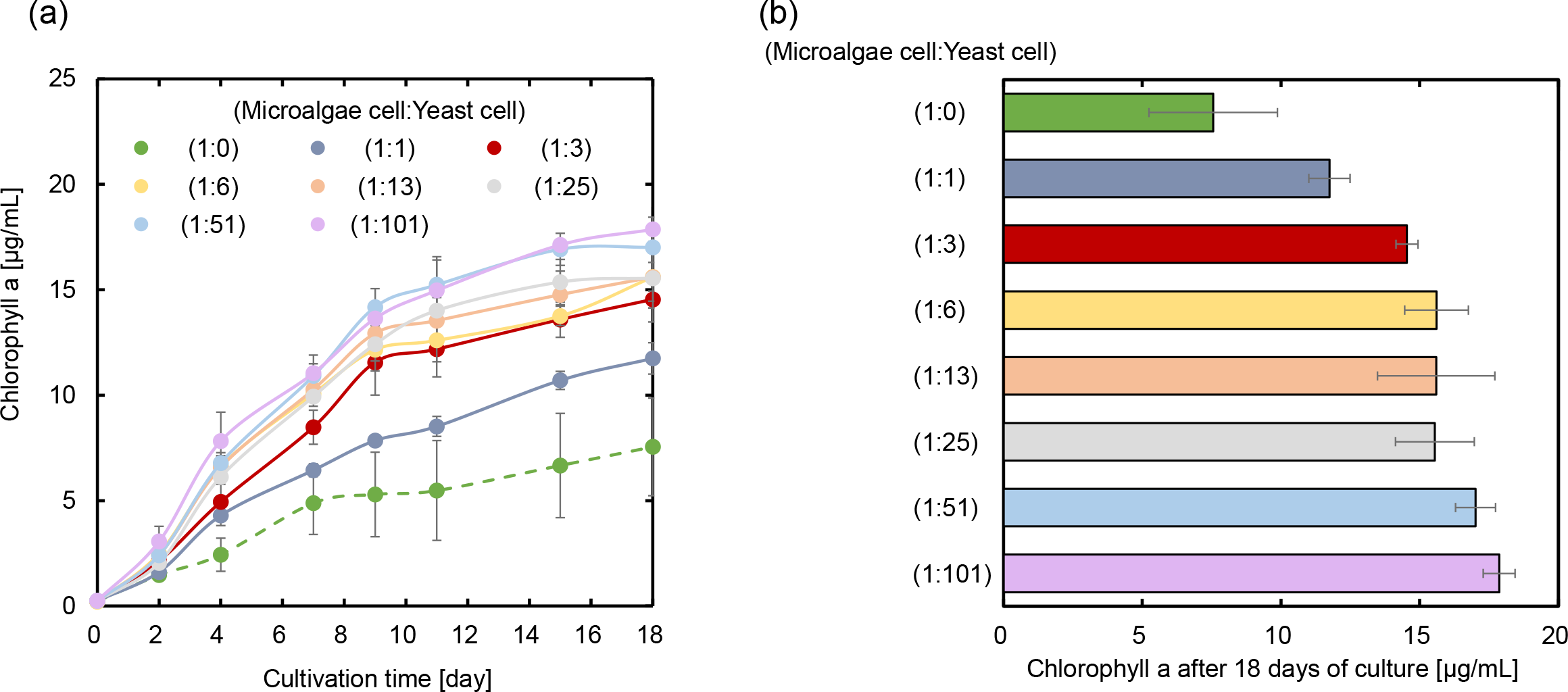
(a) Time course of chlorophyll concentration and (b) chlorophyll concentration after 18 d of culture. Three biological replicates were performed and all data reported as mean ± standard deviation.

### 3.2. Comparison of gene expression levels during co-culture and monoculture

Transcriptome analysis was performed to elucidate the cause of the increase in the cell concentration of green algae owing to co-cultivation. Total RNA was extracted from the cells after 4 d of culture under co-culture conditions (ratio of initial green algae cells to yeast cells was 1:3), which improved the growth of green algae the most, and transcriptome analysis was performed. For comparison, transcriptome analysis was also performed after 4 d of monoculture at each initial cell concentration.

The -log_10_ (p-value), which indicates statistical significance, and log_2_ (fold change), which indicates changes in gene expression levels, of each gene in the co-culture were calculated and are shown as volcano plots (Fig. 4). Genes with high log_10_ (p-value) were considered reliable, and genes with high log_2_ (fold change) showed higher transcription levels in the co-culture than in the monoculture. DEGs in the co-culture were screened using the composite criteria of a p-value less than 0.01 and at least a 2-fold change in expression (which means log_2_ [fold change] is greater than or equal to 1.0, or less than or equal to -1.0).

**Fig. 4.**
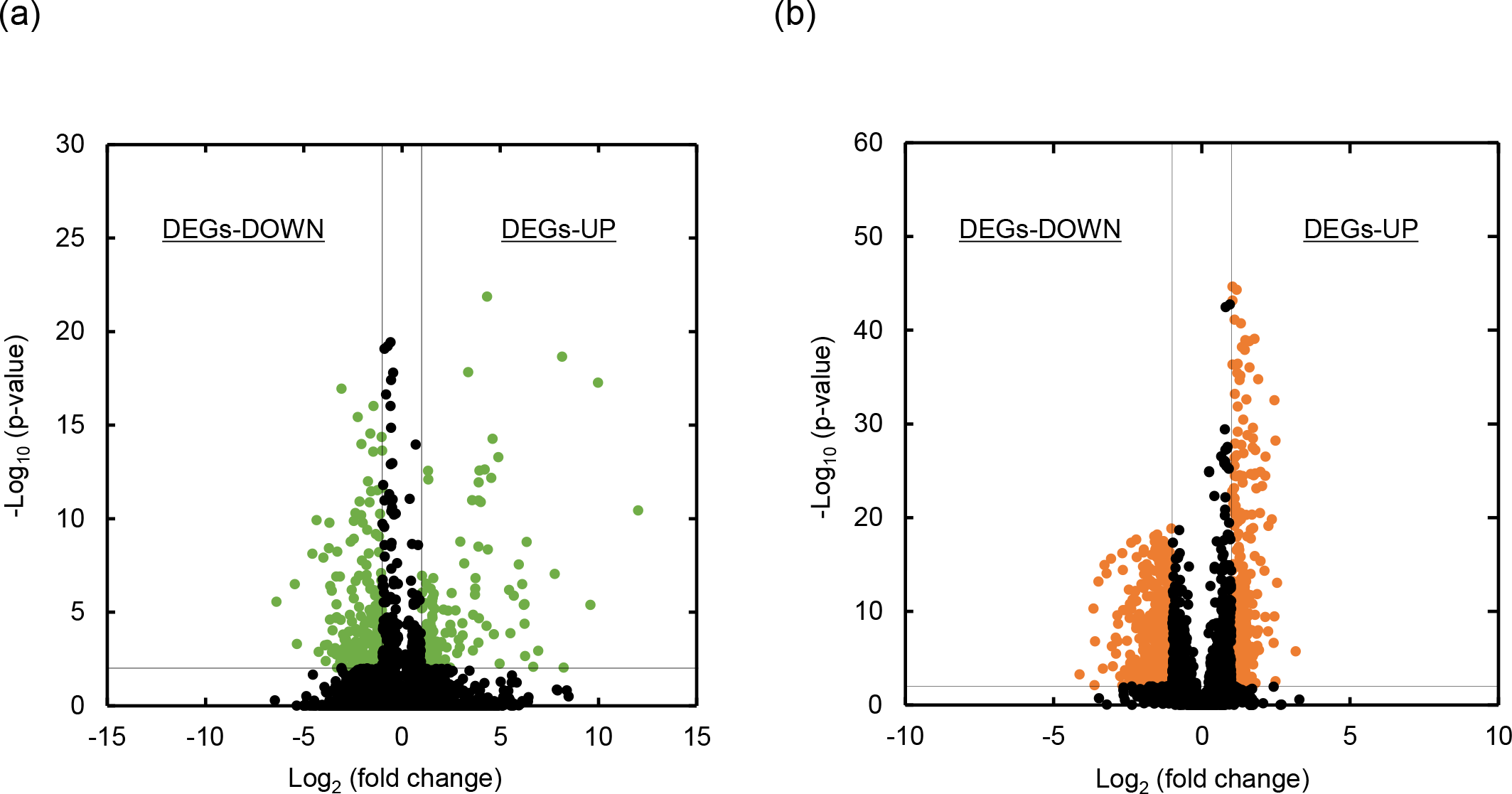
Volcano plot for genes (a) in green algae and (b) in yeast. Vertical lines represent fold changes of ±2.0 (log_2_ fold change = ±1.0). The horizontal line represents a p-value of 0.01 (-log_10_ p-value was 2.0). Green and orange points represent DEGs in green algae and yeast, respectively. Black plot points represent non-DEGs (i.e., genes whose transcription levels were unchanged).

Transcriptome analysis identified 363 DEGs, including 150 upregulated and 213 downregulated genes in green algae, and 815 DEGs, including 370 upregulated and 445 downregulated genes in yeast (Supplementary Tables 1–4). Among the identified DEGs, the top 25 high -log10 (p-values) up- and downregulated genes with known function in green algae and yeast, respectively, are summarized in Tables 1–4. In green algae, genes whose expression was increased through co-culture included CHLRE_06g284100v5 (responsible for ammonium transport), whereas genes whose expression was decreased included *CAH1* (responsible for CO_2_ enrichment), *RBCS1* (responsible for the photosynthetic dark reaction), and *PGK1* (responsible for glycogenesis). In contrast, in yeast, genes whose expression was increased through co-culture included *TDH3* (responsible for glycolysis) and *DDR2* (responsible for the stress response), whereas genes whose expression was decreased included *BAP3* (responsible for amino acid transport).

**Table 1.** Top 25 green algae genes with -Log10 (p-value) whose expression was upregulated via co-cultivation.

|  | Gene<sup>a</sup> or locus tag<sup>b</sup> | Log<sub>2</sub> (fold change) | -Log<sub>10</sub> (p-value) |
| --- | --- | --- | --- |
| 1 | CHLRE_13g590750v5 | 4.33 | 21.87 |
| 2 | CHLRE_09g413475v5 | 8.15 | 18.66 |
| 3 | CHLRE_16g651750v5 | 3.38 | 17.83 |
| 4 | CHLRE_07g350300v5 | 9.98 | 17.26 |
| 5 | CHLRE_01g017701v5 | 4.62 | 14.27 |
| 6 | CHLRE_06g284100v5 | 4.91 | 13.28 |
| 7 | CHLRE_13g590800v5 | 4.22 | 12.63 |
| 8 | CHLRE_01g054500v5 | 1.33 | 12.56 |
| 9 | CHLRE_08g377450v5 | 4.55 | 12.17 |
| 10 | CHLRE_08g372100v5 | 1.35 | 12.09 |
| 11 | CHLRE_13g591200v5 | 3.91 | 11.93 |
| 12 | CHLRE_01g026100v5 | 3.58 | 10.98 |
| 13 | CHLRE_01g012150v5 | 3.91 | 10.97 |
| 14 | CHLRE_16g652150v5 | 4.02 | 10.89 |
| 15 | CHLRE_01g062172v5 | 2.97 | 8.77 |
| 16 | CHLRE_06g284050v5 | 3.90 | 8.50 |
| 17 | CHLRE_16g687700v5 | 1.02 | 6.94 |
| 18 | CHLRE_13g591150v5 | 3.75 | 6.82 |
| 19 | CHLRE_06g250902v5 | 1.60 | 6.81 |
| 20 | CHLRE_12g527000v5 | 1.24 | 6.52 |
| 21 | CH3-II | 3.73 | 6.26 |
| 22 | CHLRE_17g714200v5 | 1.04 | 6.04 |
| 23 | CHLRE_16g675861v5 | 2.53 | 6.02 |
| 24 | CHLRE_17g726850v5 | 1.63 | 5.97 |
| 25 | CHLRE_13g567450v5 | 5.70 | 5.88 |

<sup>a</sup>UniProt (https://www.uniprot.org/)
<sup>b</sup>Gramene (https://www.gramene.org/)

## 4. Discussions

To date, there have been few examples of co-cultures of the green alga *C. reinhardtii* and yeast *S. cerevisiae*. In this study, we determined the culture conditions under which the co-cultivation of *C. reinhardtii* and *S. cerevisiae* enhanced the growth potential of *C. reinhardtii* (Fig. 1). Transcriptome analysis was also performed to identify genes whose expression was altered through co-culture with *C. reinhardtii* and *S. cerevisiae* (Fig. 4).

When green algae and yeast were co-cultured at a specific initial cell concentration ratio, the growth potential of green algae increased compared to that when they were cultured alone (Fig. 1). The growth potential of green algae was higher when the green algae to yeast ratio was 1:3 than when it was 1:1. However, when the initial yeast cell concentration was increased above 1:3 (algae cells:yeast cells), the growth potential of green algae tended to decrease with each increase in the initial yeast cell concentration. Previous studies have reported that yeasts decompose organic nitrogen compounds to produce ammonia and other compounds [17]. In general, green algae cannot utilize organic nitrogen compounds but can utilize ammonium salts and other inorganic nitrogen compounds. Therefore, it is possible that the yeast used in this study assisted the growth of green algae by converting organic nitrogen compounds into ammonium salts and other compounds in the culture medium. However, previous studies have reported that in co-cultures, when one strain grows exclusively, it inhibits the growth of other microorganisms [10]. Therefore, it is possible that the growth of green algae was inhibited when the initial yeast cell concentration was too high.

Co-culture with green algae resulted in lower yeast cell concentrations (Fig. 2). In yeast monoculture, various forms of nitrogen sources in the medium can be taken up by cells, and ammonia and glutamate can be produced and used. However, in the co-culture system, ammonia and glutamate produced by the yeast are also transferred to the green algae, which is thought to reduce the efficiency of nitrogen utilization by the yeast. This may lead to a decrease in the growth potential of the yeast in the co-culture system.

The chlorophyll concentration was higher in the co-culture than in the green algae monoculture (Fig. 3). Previous studies have reported that co-cultivation increases the chlorophyll concentration [11]. This phenomenon is thought to occur because light is shielded by the presence of heterotrophic microorganisms during co-cultivation, which suppresses photosynthesis, thus promoting chlorophyll production and compensating for the light energy required for photosynthesis [11]. In the present study, chlorophyll concentrations were particularly high at higher initial yeast inoculum concentrations, suggesting that, as in previous studies, light blocking by heterotrophic microorganisms is a factor in the increase in chlorophyll concentration.

Transcriptome analysis identified numerous green algae and yeast genes whose expression varied as a result of co-cultivation (Fig. 4, Tables 1–4). However, many of the genes, especially in green algae, had no identified function, and it was difficult to associate many of these genes with the increased growth potential of green algae owing to co-cultivation. Among the upregulated genes in green algae, CHLRE_06g284100v5 was related to ammonium transport (Table 1). In addition, the expression of *PGK1*, a glycogenesis-related gene that is upregulated during nitrogen deprivation, was downregulated [18] (Table 2). These results support the hypothesis that organic nitrogen compounds are degraded by yeast, increasing the amount of ammonium salts that are readily available to algae.

**Table 2.** Top 25 green algae genes with -Log10 (p-value) whose expression was downregulated via co-cultivation.

|  | Gene<sup>a</sup> or locus tag<sup>b</sup> | Log<sub>2</sub> (fold change) | -Log<sub>10</sub> (p-value) |
| --- | --- | --- | --- |
| 1 | CHLRE_06g303200v5 | -3.08 | 16.94 |
| 2 | CHLRE_09g392319v5 | -1.44 | 16.02 |
| 3 | CHLRE_01g030200v5 | -1.61 | 14.55 |
| 4 | CHLRE_06g259850v5 | -1.03 | 14.36 |
| 5 | CHLRE_02g079450v5 | -2.06 | 13.99 |
| 6 | CAH1 | -1.00 | 13.64 |
| 7 | CHLRE_10g428850v5 | -1.46 | 13.57 |
| 8 | RBCS1 | -1.00 | 11.64 |
| 9 | CHLRE_03g156950v5 | -1.24 | 11.51 |
| 10 | CHLRE_06g257450v5 | -1.57 | 11.46 |
| 11 | CHLRE_12g546400v5 | -1.64 | 10.87 |
| 12 | CHLRE_02g141926v5 | -2.37 | 10.29 |
| 13 | LC7 | -1.11 | 10.26 |
| 14 | CHLRE_03g206450v5 | -1.99 | 9.77 |
| 15 | COP4 | -1.76 | 9.40 |
| 16 | CHLRE_12g494800v5 | -1.32 | 9.18 |
| 17 | CHLRE_10g431800v5 | -2.61 | 8.77 |
| 18 | CHLRE_17g746997v5 | -1.17 | 8.20 |
| 19 | CHLRE_01g004250v5 | -1.68 | 8.13 |
| 20 | CHLRE_02g105950v5 | -1.81 | 7.52 |
| 21 | CHLRE_10g420700v5 | -1.06 | 7.08 |
| 22 | CHLRE_12g508900v5 | -1.79 | 6.98 |
| 23 | CHLRE_06g283950v5 | -2.18 | 6.95 |
| 24 | CHLRE_01g055400v5 | -3.15 | 6.90 |
| 25 | PGK1 | -1.04 | 6.69 |

<sup>a</sup>UniProt (https://www.uniprot.org/)
<sup>b</sup>Gramene (https://www.gramene.org/)

In contrast, the upregulated genes in yeast included *TDH3*, a gene whose expression increased during starvation [19] (Table 3). In addition, the downregulated yeast genes included *BAP3*, a gene whose expression decreased when amino acids were depleted [20] (Table 4). These results support the hypothesis that ammonia and glutamate produced by yeast in the co-culture system are transferred to green algae, thereby reducing the efficiency of nitrogen utilization by the yeast.

**Table 3.** Top 25 yeast genes with -Log10 (p-value) whose expression was upregulated via co-cultivation.

|  | Gene<sup>a</sup> or locus tag<sup>b</sup> | Log<sub>2</sub> (fold change) | -Log<sub>10</sub> (p-value) |
| --- | --- | --- | --- |
| 1 | TDH3 | 1.04 | 44.65 |
| 2 | QCR7 | 1.18 | 44.32 |
| 3 | RPL5 | 1.04 | 43.17 |
| 4 | TMA19 | 1.11 | 41.12 |
| 5 | IDI1 | 1.32 | 40.72 |
| 6 | RPC10 | 1.78 | 39.08 |
| 7 | RPP2A | 1.47 | 38.93 |
| 8 | TIM10 | 1.60 | 38.81 |
| 9 | DDR2 | 1.36 | 38.21 |
| 10 | NDE1 | 1.45 | 37.89 |
| 11 | RPL16B | 1.21 | 36.4 |
| 12 | DBP2 | 1.04 | 36.33 |
| 13 | ACC1 | 1.61 | 36.02 |
| 14 | PMA1 | 1.2 | 35.42 |
| 15 | HXK1 | 1.3 | 35.15 |
| 16 | TDH1 | 1.9 | 34.77 |
| 17 | RPS26B | 1.28 | 34.68 |
| 18 | FSH1 | 1.12 | 33.2 |
| 19 | ATP7 | 1.51 | 32.6 |
| 20 | ADE17 | 2.45 | 32.52 |
| 21 | SAH1 | 1.21 | 31.86 |
| 22 | ITR1 | 1.4 | 30.46 |
| 23 | ARO4 | 1.73 | 29.59 |
| 24 | RPL33A | 1.21 | 29.17 |
| 25 | PST1 | 1.54 | 28.81 |

<sup>a</sup>UniProt (https://www.uniprot.org/)
<sup>b</sup>*Saccharomyces* Genome Database (https://www.yeastgenome.org/)

**Table 4.** Top 25 yeast genes with -Log10 (p-value) whose expression was downregulated via co-cultivation.

|  | Gene<sup>a</sup> or locus tag<sup>b</sup> | Log<sub>2</sub> (fold change) | -Log<sub>10</sub> (p-value) |
| --- | --- | --- | --- |
| 1 | SAC6 | -1.02 | 18.84 |
| 2 | YDL012C | -1.60 | 17.98 |
| 3 | BAP3 | -1.51 | 17.92 |
| 4 | YLR108C | -2.21 | 17.66 |
| 5 | RSN1 | -1.29 | 17.47 |
| 6 | ARK1 | -1.58 | 17.43 |
| 7 | ASE1 | -1.42 | 17.43 |
| 8 | TVP18 | -1.56 | 17.41 |
| 9 | YAR029W | -2.38 | 17.32 |
| 10 | PRC1 | -1.15 | 16.96 |
| 11 | FET5 | -1.54 | 16.76 |
| 12 | YDR210W | -1.95 | 16.60 |
| 13 | NUM1 | -1.56 | 16.60 |
| 14 | YHR214C-C | -1.41 | 16.49 |
| 15 | PIL1 | -1.35 | 16.31 |
| 16 | FLC2 | -1.08 | 16.31 |
| 17 | OYE3 | -1.68 | 16.02 |
| 18 | YOL103W-A | -1.41 | 15.81 |
| 19 | GAT2 | -2.19 | 15.78 |
| 20 | THP3 | -1.51 | 15.75 |
| 21 | FMP45 | -1.24 | 15.63 |
| 22 | tP(UGG)Q | -3.06 | 15.60 |
| 23 | HUR1 | -1.43 | 15.41 |
| 24 | SSD1 | -1.43 | 15.20 |
| 25 | PMU1 | -1.86 | 15.03 |

<sup>a</sup>UniProt (https://www.uniprot.org/)
<sup>b</sup>*Saccharomyces* Genome Database (https://www.yeastgenome.org/)

Among the downregulated genes in green algae was *CAH1*, which is related to the CO_2_ enrichment mechanism (Table 2). Green algae concentrate and fix CO_2_ by actively incorporating CO_2_ and HCO_3-_ into the cell, thereby maximizing photosynthetic efficiency under CO_2_ limited conditions (CO_2_ concentrating mechanism) [21]. Previous studies have reported that electron transport around photosystem I (PSI) is involved in the uptake of inorganic carbon and that under low CO_2_ concentrations, an excess amount of energy is sent from PSI to drive the inorganic carbon pump, resulting in a high PSI/photosystem II (PSII) ratio. The energy available for the Calvin cycle decreases when the CO_2_ enrichment mechanism is active, resulting in a decrease in the growth rate and CO_2_ fixation efficiency of green algae [22,23]. Therefore, it is likely that the CO_2_ produced by yeast in the co-culture system increased the concentration of inorganic carbon in the culture medium, resulting in a decrease in gene transcription of *CAH1*. This could also be attributed to the increased growth potential of green algae owing to co-cultivation. Furthermore, *DDR2*, a gene associated with oxidative stress, was included among the upregulated genes in yeast (Table 3). *DDR2* encodes a protein that removes damaged proteins and toxic oxygen species when oxidative stress is applied to yeast [24]. In the co-culture system used in this study, *DDR2* transcription increased as a result of oxidative stress in yeast due to O_2_ produced by green algae. Therefore, these results support the exchange of O_2_ and CO_2_ between green algae and yeast in co-culture. However, there have been few reports showing direct evidence of O_2_ and CO_2_ exchange between microalgae and heterotrophic microorganisms. Thus, further studies are required.

The gene *RBCS1*, related to the photosynthetic dark reaction, was also among the downregulated genes in green algae (Table 2). In the photosynthetic dark reaction, the ATP and NADPH _2_^+^ generated in the light reaction are used to produce sugars from CO_2_. *RBCS1* expression is induced by light [25]. Therefore, the lower gene expression of *RBCS1* supports the previous hypothesis that the presence of yeast during co-culture blocks light and suppresses photosynthesis, thus promoting chlorophyll production to compensate for the light energy required by green algae.

## 5. Conclusions

In this study, we found that the growth potential of *C. reinhardtii* was enhanced by co-culturing the green algae *C. reinhardtii* with the yeast *S. cerevisiae*. We also performed transcriptome analysis of *C. reinhardtii* and *S. cerevisiae* under culture conditions that increased the growth potential of *C. reinhardtii* and discussed the cause of the increased growth potential of *C. reinhardtii*. Although further studies are needed to elucidate the full impact of microbial interactions in *C. reinhardtii* and *S. cerevisiae* co-cultures, the results of the transcriptome analysis in this study represent an important first step toward this goal. Further research on microalgae and heterotrophic co-cultures is expected to lead to the use of microalgae co-cultures for biological CO_2_ fixation and production of food and feed from atmospheric CO_2_.

