## Supplementary material for "Improvement of cell growth in green algae *Chlamydomonas reinhardtii* through co-cultivation with yeast *Saccharomyces cerevisiae*"

**Supplementary Table 1 Genes in green algae upregulated via co-cultivation**

| Gene[a] or locus tag[b] | $Log_2$ (fold change) | $-Log_{10}$ (p-value) |
|---|---|---|
| CHLRE_13g590750v5 | 4.33 | 21.87 |
| CHLRE_09g413475v5 | 8.15 | 18.66 |
| CHLRE_16g651750v5 | 3.38 | 17.83 |
| CHLRE_07g350300v5 | 9.98 | 17.26 |
| CHLRE_01g017701v5 | 4.62 | 14.27 |
| CHLRE_06g284100v5 | 4.91 | 13.28 |
| CHLRE_13g590800v5 | 4.22 | 12.63 |
| CHLRE_05g235355v5 | 3.95 | 12.57 |
| CHLRE_01g054500v5 | 1.33 | 12.56 |
| CHLRE_08g377450v5 | 4.55 | 12.17 |
| CHLRE_08g372100v5 | 1.35 | 12.09 |
| CHLRE_13g591200v5 | 3.91 | 11.93 |
| CHLRE_01g026100v5 | 3.58 | 10.98 |
| CHLRE_01g012150v5 | 3.91 | 10.97 |
| CHLRE_16g652150v5 | 4.02 | 10.89 |
| CHLRE_09g390838v5 | 12.02 | 10.44 |
| CHLRE_01g062172v5 | 2.97 | 8.77 |
| CHLRE_17g734725v5 | 6.35 | 8.75 |
| CHLRE_06g284050v5 | 3.90 | 8.50 |
| CHLRE_13g605300v5 | 4.38 | 8.35 |
| CHLRE_13g590150v5 | 3.17 | 7.60 |
| CHLRE_06g282826v5 | 5.95 | 7.54 |
| CHLRE_02g097600v5 | 7.78 | 7.04 |
| CHLRE_16g687700v5 | 1.02 | 6.94 |
| CHLRE_13g591150v5 | 3.75 | 6.82 |
| CHLRE_06g250902v5 | 1.60 | 6.81 |
| CHLRE_12g527000v5 | 1.24 | 6.52 |
| CHLRE_07g327317v5 | 6.12 | 6.50 |
| CH3-II | 3.73 | 6.26 |
| CHLRE_01g015500v5 | 1.46 | 6.26 |
| CHLRE_07g344260v5 | 5.45 | 6.19 |
| CHLRE_17g714200v5 | 1.04 | 6.04 |
| CHLRE_16g675861v5 | 2.53 | 6.02 |
| CHLRE_17g726850v5 | 1.63 | 5.97 |
| CHLRE_09g389245v5 | 3.72 | 5.94 |

| | | |
|---|---|---|
| CHLRE_13g567450v5 | 5.70 | 5.88 |
| CHLRE_12g500550v5 | 1.57 | 5.77 |
| CHLRE_02g095076v5 | 1.53 | 5.70 |
| CHLRE_07g347400v5 | 1.54 | 5.60 |
| CHLRE_01g050950v5 | 1.16 | 5.48 |
| CHLRE_09g392803v5 | 6.24 | 5.43 |
| CHLRE_06g278273v5 | 9.59 | 5.38 |
| CHLRE_16g649950v5 | 6.21 | 5.38 |
| CHLRE_13g574900v5 | 1.56 | 5.33 |
| CHLRE_10g435350v5 | 1.05 | 5.24 |
| CHLRE_08g385350v5 | 1.01 | 5.21 |
| CHLRE_04g217300v5 | 1.66 | 5.17 |
| CHLRE_04g223300v5 | 2.12 | 5.13 |
| CHLRE_06g295150v5 | 2.73 | 5.09 |
| CHLRE_17g746547v5 | 2.34 | 5.08 |
| CHLRE_15g643384v5 | 3.64 | 4.85 |
| CHLRE_02g112750v5 | 1.54 | 4.82 |
| CHLRE_11g467649v5 | 1.56 | 4.70 |
| CHLRE_03g167778v5 | 3.92 | 4.67 |
| CHLRE_08g368026v5 | 3.08 | 4.41 |
| CHLRE_17g701050v5 | 1.45 | 4.40 |
| CHLRE_05g233602v5 | 6.24 | 4.37 |
| CHLRE_12g529600v5 | 1.26 | 4.36 |
| CHLRE_06g266800v5 | 2.46 | 4.32 |
| CHLRE_09g392951v5 | 4.31 | 4.27 |
| CHLRE_07g347300v5 | 1.50 | 4.16 |
| CHLRE_04g224600v5 | 1.41 | 4.12 |
| CHLRE_08g378800v5 | 1.30 | 4.08 |
| CHLRE_10g453050v5 | 1.22 | 4.08 |
| CHLRE_01g029950v5 | 1.23 | 4.03 |
| CHLRE_13g588650v5 | 1.33 | 4.01 |
| CHLRE_07g350750v5 | 2.51 | 3.98 |
| CHLRE_15g634650v5 | 1.60 | 3.97 |
| CHLRE_14g616826v5 | 2.53 | 3.92 |
| CHLRE_01g041100v5 | 2.22 | 3.89 |
| CHLRE_01g007901v5 | 1.54 | 3.88 |
| CHLRE_13g591073v5 | 5.52 | 3.87 |

| | | |
|---|---|---|
| CHLRE_12g536683v5 | 4.68 | 3.81 |
| CHLRE_12g534800v5 | 1.47 | 3.80 |
| CHLRE_12g520000v5 | 3.10 | 3.76 |
| CHLRE_02g082852v5 | 1.48 | 3.67 |
| CHLRE_04g215600v5 | 1.49 | 3.58 |
| CHLRE_05g238250v5 | 1.09 | 3.54 |
| CHLRE_17g718050v5 | 2.90 | 3.52 |
| CHLRE_01g004800v5 | 1.03 | 3.52 |
| CHLRE_17g714150v5 | 1.77 | 3.42 |
| CHLRE_11g467547v5 | 1.14 | 3.42 |
| CHLRE_11g468600v5 | 1.53 | 3.40 |
| CHLRE_13g563600v5 | 2.01 | 3.39 |
| CHLRE_09g392914v5 | 3.90 | 3.37 |
| CHLRE_17g697406v5 | 1.32 | 3.30 |
| CHLRE_07g330300v5 | 1.11 | 3.29 |
| CHLRE_02g091950v5 | 1.66 | 3.28 |
| CHLRE_03g188400v5 | 1.40 | 3.25 |
| CHLRE_09g396950v5 | 1.19 | 3.18 |
| CHLRE_07g317450v5 | 1.92 | 3.13 |
| CHLRE_08g379400v5 | 1.79 | 3.13 |
| CHLRE_09g389503v5 | 2.95 | 3.12 |
| CHLRE_11g467644v5 | 2.28 | 3.11 |
| CHLRE_14g620500v5 | 1.28 | 3.08 |
| CHLRE_09g398050v5 | 1.79 | 3.03 |
| CHLRE_07g329750v5 | 1.18 | 3.01 |
| CHLRE_16g650250v5 | 3.61 | 2.95 |
| CHLRE_07g353450v5 | 1.47 | 2.95 |
| CHLRE_09g399030v5 | 1.15 | 2.94 |
| CHLRE_06g271550v5 | 6.94 | 2.93 |
| CHLRE_17g733100v5 | 2.81 | 2.81 |
| CHLRE_04g217971v5 | 1.15 | 2.80 |
| CHLRE_16g668300v5 | 1.78 | 2.79 |
| CHLRE_11g477550v5 | 1.10 | 2.79 |
| CHLRE_02g099550v5 | 1.31 | 2.77 |
| CHLRE_03g160400v5 | 1.02 | 2.74 |
| CHLRE_12g519900v5 | 1.29 | 2.71 |
| CHLRE_01g053150v5 | 1.75 | 2.69 |

| | | |
|---|---|---|
| CHLRE_07g332800v5 | 1.16 | 2.69 |
| CHLRE_16g691800v5 | 1.01 | 2.68 |
| CHLRE_08g360150v5 | 6.27 | 2.65 |
| CHLRE_12g487200v5 | 1.09 | 2.65 |
| CHLRE_09g409650v5 | 1.27 | 2.64 |
| CHLRE_09g387208v5 | 1.36 | 2.57 |
| CHLRE_12g501600v5 | 1.02 | 2.50 |
| CHLRE_09g397450v5 | 2.20 | 2.47 |
| CHLRE_12g558300v5 | 1.63 | 2.45 |
| CHLRE_12g558352v5 | 1.05 | 2.44 |
| CHLRE_12g496400v5 | 1.63 | 2.41 |
| CHLRE_08g373366v5 | 1.15 | 2.40 |
| CHLRE_03g144847v5 | 1.95 | 2.38 |
| CHLRE_10g461400v5 | 1.66 | 2.36 |
| CHLRE_08g384150v5 | 1.50 | 2.36 |
| CHLRE_02g142867v5 | 1.90 | 2.35 |
| CHLRE_04g224800v5 | 1.87 | 2.34 |
| CHLRE_11g474900v5 | 1.07 | 2.31 |
| CHLRE_04g213152v5 | 1.47 | 2.30 |
| CHLRE_13g589450v5 | 1.33 | 2.30 |
| CHLRE_17g721400v5 | 1.49 | 2.28 |
| CHLRE_05g232200v5 | 1.15 | 2.27 |
| CHLRE_03g155250v5 | 1.02 | 2.27 |
| CHLRE_15g642865v5 | 4.98 | 2.25 |
| CHLRE_06g250100v5 | 1.51 | 2.25 |
| CHLRE_05g248200v5 | 1.06 | 2.25 |
| CHLRE_05g237950v5 | 1.08 | 2.21 |
| CHLRE_08g358250v5 | 1.43 | 2.20 |
| CHLRE_06g288700v5 | 1.13 | 2.20 |
| CHLRE_01g005150v5 | 1.19 | 2.18 |
| CHLRE_06g287850v5 | 1.58 | 2.17 |
| CHLRE_01g046650v5 | 1.57 | 2.15 |
| CHLRE_05g234660v5 | 1.06 | 2.10 |
| CHLRE_02g142987v5 | 6.68 | 2.08 |
| CHLRE_09g399350v5 | 1.08 | 2.06 |
| CHLRE_09g403150v5 | 1.28 | 2.05 |
| CHLRE_06g272600v5 | 8.23 | 2.04 |

| | | |
|---|---|---|
| CHLRE_12g553552v5 | 2.45 | 2.03 |
| CHLRE_12g515500v5 | 1.58 | 2.02 |
| CHLRE_07g347750v5 | 1.38 | 2.01 |
| CHLRE_01g014650v5 | 1.40 | 2.00 |

[a]UniProt (https://www.uniprot.org/)

[b]Gramene (https://www.gramene.org/)

**Supplementary Table 2 Genes in green algae downregulated via co-cultivation**

| Gene[a] or locus tag[b] | $Log_2$ (fold change) | $-Log_{10}$ (p-value) |
|---|---|---|
| CHLRE_06g303200v5 | -3.08 | 16.94 |
| CHLRE_09g392319v5 | -1.44 | 16.02 |
| CHLRE_08g367550v5 | -2.24 | 15.43 |
| CHLRE_01g030200v5 | -1.61 | 14.55 |
| CHLRE_06g259850v5 | -1.03 | 14.36 |
| CHLRE_02g079450v5 | -2.06 | 13.99 |
| CAH1 | -1.00 | 13.64 |
| CHLRE_10g428850v5 | -1.46 | 13.57 |
| CHLRE_12g495550v5 | -1.72 | 11.99 |
| RBCS1 | -1.00 | 11.64 |
| CHLRE_03g156950v5 | -1.24 | 11.51 |
| CHLRE_06g257450v5 | -1.57 | 11.46 |
| CHLRE_03g157450v5 | -2.15 | 10.91 |
| CHLRE_12g546400v5 | -1.64 | 10.87 |
| CHLRE_02g141926v5 | -2.37 | 10.29 |
| LC7 | -1.11 | 10.26 |
| CHLRE_02g085300v5 | -2.07 | 10.18 |
| CHLRE_10g442050v5 | -4.34 | 9.92 |
| CHLRE_17g725400v5 | -2.44 | 9.88 |
| CHLRE_11g468359v5 | -3.69 | 9.78 |
| CHLRE_03g206450v5 | -1.99 | 9.77 |
| COP4 | -1.76 | 9.40 |
| CHLRE_12g494800v5 | -1.32 | 9.18 |
| CHLRE_06g251000v5 | -1.17 | 9.05 |
| CHLRE_02g074737v5 | -2.45 | 8.92 |
| CHLRE_10g431800v5 | -2.61 | 8.77 |
| CHLRE_17g710300v5 | -3.71 | 8.42 |
| CHLRE_10g448900v5 | -3.29 | 8.24 |
| CHLRE_17g746997v5 | -1.17 | 8.20 |
| CHLRE_01g004250v5 | -1.68 | 8.13 |
| CHLRE_10g463370v5 | -4.55 | 8.12 |
| CHLRE_09g409901v5 | -4.00 | 7.91 |
| CHLRE_04g216250v5 | -2.03 | 7.75 |
| CHLRE_02g105950v5 | -1.81 | 7.52 |
| CHLRE_10g420700v5 | -1.06 | 7.08 |

| | | |
|---|---|---|
| CHLRE_12g508900v5 | -1.79 | 6.98 |
| CHLRE_06g283950v5 | -2.18 | 6.95 |
| CHLRE_07g328750v5 | -1.83 | 6.94 |
| CHLRE_01g055400v5 | -3.15 | 6.90 |
| CHLRE_02g095150v5 | -3.34 | 6.90 |
| PGK1 | -1.04 | 6.69 |
| CHLRE_06g251600v5 | -2.35 | 6.66 |
| CHLRE_10g427300v5 | -1.77 | 6.64 |
| CHLRE_16g683150v5 | -1.66 | 6.55 |
| CHLRE_05g238600v5 | -5.45 | 6.49 |
| CHLRE_15g642800v5 | -3.65 | 6.39 |
| CHLRE_01g005450v5 | -1.61 | 6.24 |
| CHLRE_09g395213v5 | -2.55 | 6.18 |
| CHLRE_07g316000v5 | -1.42 | 6.17 |
| CHLRE_09g399250v5 | -1.25 | 6.15 |
| CHLRE_17g705500v5 | -3.57 | 6.14 |
| CHLRE_16g670800v5 | -1.24 | 5.90 |
| CHLRE_16g655200v5 | -1.36 | 5.89 |
| CHLRE_04g216203v5 | -2.45 | 5.73 |
| CHLRE_05g243472v5 | -6.38 | 5.56 |
| CHLRE_12g499800v5 | -1.20 | 5.46 |
| CHLRE_16g693500v5 | -3.33 | 5.42 |
| CHLRE_02g141166v5 | -1.34 | 5.35 |
| CHLRE_10g451752v5 | -2.18 | 5.28 |
| CHLRE_10g432850v5 | -1.18 | 5.27 |
| CHLRE_12g486250v5 | -1.18 | 5.19 |
| CHLRE_17g714250v5 | -1.64 | 5.12 |
| CHLRE_16g657650v5 | -1.62 | 5.11 |
| CHLRE_08g373359v5 | -1.02 | 5.10 |
| CHLRE_03g200950v5 | -1.04 | 4.99 |
| CHLRE_02g089950v5 | -1.35 | 4.94 |
| CHLRE_17g721250v5 | -1.55 | 4.90 |
| CHLRE_07g327400v5 | -1.20 | 4.88 |
| CHLRE_06g252743v5 | -2.64 | 4.85 |
| CHLRE_03g201950v5 | -2.08 | 4.79 |
| CHLRE_06g276200v5 | -1.21 | 4.78 |
| CHLRE_09g398900v5 | -3.32 | 4.73 |

| | | |
|---|---|---|
| CHLRE_16g689311v5 | -3.36 | 4.68 |
| CHLRE_01g051250v5 | -1.36 | 4.66 |
| CHLRE_04g228950v5 | -1.57 | 4.65 |
| CHLRE_14g615634v5 | -3.66 | 4.60 |
| CHLRE_07g330750v5 | -3.12 | 4.59 |
| CHLRE_17g734757v5 | -1.38 | 4.56 |
| CHLRE_09g410600v5 | -1.01 | 4.54 |
| CHLRE_07g353325v5 | -1.82 | 4.52 |
| CHLRE_06g263200v5 | -2.54 | 4.49 |
| CHLRE_02g106450v5 | -1.52 | 4.41 |
| CHLRE_04g217900v5 | -1.21 | 4.35 |
| CHLRE_09g389150v5 | -1.90 | 4.33 |
| CHLRE_02g095084v5 | -1.13 | 4.31 |
| CHLRE_02g109950v5 | -1.07 | 4.29 |
| CHLRE_13g565800v5 | -1.07 | 4.29 |
| CHLRE_17g700425v5 | -1.20 | 4.28 |
| CHLRE_09g408200v5 | -1.18 | 4.27 |
| CHLRE_17g700133v5 | -1.24 | 4.11 |
| CHLRE_06g278170v5 | -2.34 | 4.10 |
| CHLRE_10g455300v5 | -1.24 | 4.02 |
| CHLRE_11g481600v5 | -3.53 | 4.02 |
| CHLRE_02g090050v5 | -1.97 | 3.98 |
| CHLRE_05g238650v5 | -2.70 | 3.95 |
| CHLRE_06g289426v5 | -2.28 | 3.92 |
| CHLRE_08g368200v5 | -1.79 | 3.88 |
| CHLRE_02g141005v5 | -1.73 | 3.84 |
| CHLRE_17g730550v5 | -2.99 | 3.80 |
| CHLRE_12g517150v5 | -1.16 | 3.71 |
| CHLRE_12g511050v5 | -1.22 | 3.71 |
| CHLRE_16g665000v5 | -1.14 | 3.70 |
| CHLRE_16g674400v5 | -1.56 | 3.69 |
| CHLRE_01g044050v5 | -1.92 | 3.56 |
| CHLRE_10g458400v5 | -1.45 | 3.54 |
| CHLRE_13g571150v5 | -1.28 | 3.53 |
| CHLRE_01g052350v5 | -2.85 | 3.53 |
| CHLRE_01g008300v5 | -2.91 | 3.53 |
| CHLRE_06g273413v5 | -1.86 | 3.45 |

| | | |
|---|---|---|
| CHLRE_12g511200v5 | -1.62 | 3.43 |
| CHLRE_17g729450v5 | -2.34 | 3.42 |
| CHLRE_09g410800v5 | -1.80 | 3.41 |
| CHLRE_16g657250v5 | -1.53 | 3.38 |
| CHLRE_02g099950v5 | -1.31 | 3.36 |
| CHLRE_10g447100v5 | -1.13 | 3.34 |
| CHLRE_02g145231v5 | -1.32 | 3.34 |
| CHLRE_17g696700v5 | -5.34 | 3.29 |
| CHLRE_06g256450v5 | -1.87 | 3.28 |
| CHLRE_17g697850v5 | -2.62 | 3.26 |
| CHLRE_05g235850v5 | -3.79 | 3.26 |
| CHLRE_03g145267v5 | -1.48 | 3.24 |
| CHLRE_02g102050v5 | -2.44 | 3.24 |
| CHLRE_04g217963v5 | -2.85 | 3.24 |
| CHLRE_01g011150v5 | -3.91 | 3.22 |
| CHLRE_02g078550v5 | -1.30 | 3.21 |
| CHLRE_12g513650v5 | -1.47 | 3.20 |
| CHLRE_06g308050v5 | -2.36 | 3.20 |
| CHLRE_05g238311v5 | -1.39 | 3.17 |
| CHLRE_06g302305v5 | -1.10 | 3.12 |
| CHLRE_11g476850v5 | -1.34 | 3.09 |
| CHLRE_12g528850v5 | -1.04 | 3.08 |
| CHLRE_01g009950v5 | -1.41 | 3.07 |
| CHLRE_15g642976v5 | -3.39 | 3.07 |
| CHLRE_02g094450v5 | -2.54 | 3.06 |
| CHLRE_12g489001v5 | -2.74 | 3.06 |
| CHLRE_02g142650v5 | -2.21 | 3.02 |
| CHLRE_12g529100v5 | -1.18 | 3.00 |
| CHLRE_03g168500v5 | -2.78 | 3.00 |
| CHLRE_05g238687v5 | -3.02 | 3.00 |
| CHLRE_06g276400v5 | -1.22 | 2.99 |
| CHLRE_03g178350v5 | -1.26 | 2.99 |
| CHLRE_16g681850v5 | -1.34 | 2.93 |
| CHLRE_12g538950v5 | -1.46 | 2.88 |
| CHLRE_01g047950v5 | -1.15 | 2.87 |
| CHLRE_12g558700v5 | -1.41 | 2.87 |
| CHLRE_04g222402v5 | -4.23 | 2.87 |

| | | |
|---|---|---|
| CHLRE_09g392467v5 | -1.33 | 2.86 |
| CHLRE_10g442041v5 | -2.31 | 2.86 |
| CHLRE_01g012700v5 | -3.56 | 2.86 |
| CHLRE_02g074720v5 | -1.97 | 2.85 |
| CHLRE_02g143367v5 | -3.40 | 2.81 |
| CHLRE_15g643388v5 | -2.11 | 2.76 |
| CHLRE_10g433476v5 | -3.34 | 2.74 |
| CHLRE_17g738000v5 | -1.44 | 2.72 |
| CHLRE_02g085450v5 | -1.63 | 2.69 |
| CHLRE_16g649050v5 | -1.35 | 2.68 |
| CHLRE_06g278152v5 | -1.88 | 2.67 |
| CHLRE_09g391652v5 | -2.68 | 2.66 |
| CHLRE_16g670755v5 | -1.36 | 2.57 |
| CHLRE_03g197513v5 | -1.93 | 2.56 |
| CHLRE_17g733208v5 | -1.10 | 2.55 |
| CHLRE_16g657200v5 | -2.56 | 2.53 |
| CHLRE_16g670650v5 | -2.97 | 2.52 |
| CHLRE_03g151950v5 | -2.07 | 2.50 |
| CHLRE_09g413500v5 | -1.11 | 2.49 |
| CHLRE_01g000900v5 | -1.37 | 2.49 |
| CHLRE_16g671750v5 | -2.26 | 2.49 |
| CHLRE_02g116400v5 | -1.90 | 2.48 |
| CHLRE_13g568316v5 | -2.30 | 2.48 |
| CHLRE_07g341550v5 | -1.20 | 2.46 |
| CHLRE_06g260700v5 | -1.25 | 2.44 |
| CHLRE_12g535650v5 | -1.53 | 2.40 |
| CHLRE_03g159900v5 | -1.61 | 2.40 |
| CHLRE_16g693600v5 | -2.30 | 2.40 |
| CHLRE_09g389050v5 | -3.88 | 2.39 |
| CHLRE_02g142206v5 | -1.00 | 2.38 |
| CHLRE_01g033400v5 | -2.35 | 2.38 |
| CHLRE_01g025450v5 | -1.12 | 2.36 |
| CHLRE_17g722050v5 | -1.40 | 2.35 |
| CHLRE_10g433800v5 | -2.46 | 2.33 |
| CHLRE_08g365100v5 | -3.14 | 2.33 |
| CHLRE_08g385200v5 | -1.00 | 2.30 |
| CHLRE_17g724200v5 | -1.03 | 2.29 |

| | | |
|---|---|---|
| CHLRE_17g717850v5 | -2.51 | 2.29 |
| CHLRE_14g624000v5 | -2.48 | 2.28 |
| CHLRE_10g445600v5 | -1.71 | 2.26 |
| CHLRE_02g089450v5 | -1.87 | 2.24 |
| CHLRE_16g658800v5 | -2.45 | 2.24 |
| CHLRE_01g007600v5 | -1.27 | 2.21 |
| CHLRE_14g611100v5 | -2.47 | 2.21 |
| CHLRE_02g114700v5 | -1.12 | 2.20 |
| CHLRE_12g516700v5 | -1.43 | 2.18 |
| CHLRE_12g516717v5 | -1.43 | 2.18 |
| CHLRE_15g637401v5 | -2.04 | 2.18 |
| CHLRE_12g526000v5 | -3.12 | 2.17 |
| CHLRE_10g441450v5 | -1.58 | 2.16 |
| CHLRE_12g553300v5 | -1.01 | 2.15 |
| CHLRE_02g112500v5 | -2.18 | 2.13 |
| CHLRE_03g144564v5 | -1.79 | 2.12 |
| CHLRE_12g538750v5 | -1.89 | 2.10 |
| CHLRE_17g705300v5 | -2.31 | 2.10 |
| CHLRE_15g643702v5 | -1.34 | 2.09 |
| CHLRE_17g737100v5 | -1.10 | 2.08 |
| CHLRE_10g432600v5 | -1.83 | 2.07 |
| CHLRE_06g278232v5 | -2.88 | 2.06 |
| CHLRE_02g102200v5 | -1.70 | 2.05 |
| CHLRE_16g693601v5 | -2.77 | 2.05 |
| CHLRE_12g549000v5 | -3.30 | 2.04 |
| CHLRE_02g092100v5 | -1.09 | 2.03 |
| CHLRE_17g727801v5 | -2.52 | 2.03 |
| CHLRE_03g174450v5 | -1.52 | 2.02 |
| CHLRE_03g189250v5 | -1.29 | 2.01 |
| CHLRE_17g715500v5 | -2.24 | 2.01 |

[a]UniProt (https://www.uniprot.org/)

[b]Gramene (https://www.gramene.org/)

**Supplementary Table 3 Genes in yeast upregulated via co-cultivation**

| Gene[a] or locus tag[b] | $Log_2$ (fold change) | $-Log_{10}$ (p-value) |
|---|---|---|
| TDH3 | 1.04 | 44.65 |
| QCR7 | 1.18 | 44.32 |
| RPL5 | 1.04 | 43.17 |
| TMA19 | 1.11 | 41.12 |
| IDI1 | 1.32 | 40.72 |
| RPC10 | 1.78 | 39.08 |
| RPP2A | 1.47 | 38.93 |
| TIM10 | 1.60 | 38.81 |
| DDR2 | 1.36 | 38.21 |
| NDE1 | 1.45 | 37.89 |
| RPL16B | 1.21 | 36.40 |
| DBP2 | 1.04 | 36.33 |
| ACC1 | 1.61 | 36.02 |
| PMA1 | 1.20 | 35.42 |
| HXK1 | 1.30 | 35.15 |
| TDH1 | 1.90 | 34.77 |
| RPS26B | 1.28 | 34.68 |
| FSH1 | 1.12 | 33.20 |
| ATP7 | 1.51 | 32.60 |
| ADE17 | 2.45 | 32.52 |
| SAH1 | 1.21 | 31.86 |
| ITR1 | 1.40 | 30.46 |
| ARO4 | 1.73 | 29.59 |
| RPL33A | 1.21 | 29.17 |
| PST1 | 1.54 | 28.81 |
| INO1 | 1.73 | 28.45 |
| PHO3 | 2.49 | 28.20 |
| RPS10A | 1.13 | 27.89 |
| ALD5 | 1.34 | 27.75 |
| YHB1 | 1.72 | 27.49 |
| TYE7 | 1.81 | 27.21 |
| SUC2 | 1.41 | 26.86 |
| FAS1 | 1.18 | 26.64 |
| HSP150 | 2.15 | 26.49 |
| COX5A | 1.11 | 26.40 |

| | | |
|---|---|---|
| TIF11 | 1.11 | 25.54 |
| ZTA1 | 1.98 | 24.87 |
| STF2 | 1.81 | 24.71 |
| ILV3 | 1.66 | 24.66 |
| COX6 | 1.28 | 24.52 |
| YER053C-A | 1.75 | 24.50 |
| MTC7 | 2.14 | 24.47 |
| CDC33 | 1.12 | 24.47 |
| KNH1 | 1.38 | 24.43 |
| FAS2 | 1.16 | 24.40 |
| STF1 | 1.38 | 23.73 |
| TKL1 | 2.04 | 23.37 |
| RPL21A | 1.09 | 23.13 |
| CWP2 | 1.84 | 23.11 |
| KRS1 | 1.03 | 22.81 |
| ARO3 | 1.02 | 22.15 |
| RPL11A | 1.09 | 22.10 |
| RPL8B | 1.11 | 22.04 |
| RPS28B | 1.06 | 21.50 |
| ADE12 | 1.14 | 21.24 |
| RGI2 | 1.09 | 20.90 |
| MRT4 | 1.30 | 20.51 |
| SFC1 | 1.97 | 20.48 |
| AI1 | 1.13 | 20.37 |
| RKI1 | 1.69 | 20.31 |
| LHP1 | 1.50 | 20.29 |
| YPL276W | 2.36 | 19.81 |
| RPL13A | 1.32 | 19.80 |
| TIF3 | 1.22 | 19.57 |
| RPS10B | 1.05 | 19.30 |
| YCR087C-A | 2.25 | 19.11 |
| NHP2 | 1.11 | 19.10 |
| CNL1 | 1.74 | 18.88 |
| ADE1 | 1.66 | 18.71 |
| EXG1 | 1.24 | 18.27 |
| BRX1 | 1.53 | 17.98 |
| VMA7 | 1.18 | 17.92 |

| | | |
|---|---|---|
| URA7 | 1.66 | 17.76 |
| NCE103 | 1.25 | 16.88 |
| SUP45 | 1.27 | 16.55 |
| VTC1 | 1.18 | 16.52 |
| ERG20 | 1.79 | 15.89 |
| RRB1 | 1.31 | 15.82 |
| SSS1 | 1.28 | 15.71 |
| FAR1 | 1.98 | 15.36 |
| ATP4 | 1.06 | 15.02 |
| SES1 | 1.39 | 14.80 |
| ENO2 | 1.29 | 14.78 |
| RHO5 | 1.24 | 14.73 |
| FKS1 | 1.03 | 14.38 |
| CYB5 | 2.12 | 14.32 |
| QCR2 | 1.41 | 14.09 |
| ARO8 | 1.65 | 13.97 |
| YLR363W-A | 1.40 | 13.95 |
| RPL20A | 1.32 | 13.44 |
| SDH3 | 1.42 | 13.37 |
| MAK21 | 1.50 | 13.21 |
| PFK1 | 1.28 | 13.21 |
| MIN7 | 1.06 | 13.21 |
| FUR1 | 1.41 | 13.07 |
| HOM2 | 1.41 | 13.07 |
| BIO2 | 2.54 | 13.03 |
| MCR1 | 1.30 | 12.71 |
| LOC1 | 1.45 | 12.58 |
| ATP16 | 1.37 | 12.52 |
| HEM3 | 1.27 | 12.46 |
| CDC19 | 1.13 | 12.44 |
| TPM1 | 1.26 | 12.39 |
| CHO2 | 1.16 | 12.26 |
| YBR221W-A | 1.40 | 12.20 |
| ERG28 | 1.48 | 12.03 |
| ATP5 | 1.44 | 11.86 |
| NUG1 | 1.03 | 11.85 |
| MIC19 | 1.87 | 11.84 |

| | | |
|---|---|---|
| ECM1 | 1.65 | 11.72 |
| NOC2 | 1.21 | 11.69 |
| HAS1 | 1.20 | 11.48 |
| ARC19 | 1.44 | 11.08 |
| POP1 | 1.62 | 10.90 |
| PDC6 | 1.35 | 10.86 |
| YGR035W-A | 1.71 | 10.78 |
| MIS1 | 1.43 | 10.58 |
| RPA49 | 1.31 | 10.42 |
| ATP20 | 1.19 | 10.37 |
| ADE13 | 1.20 | 10.11 |
| ELF1 | 1.13 | 10.02 |
| MAK16 | 1.43 | 9.99 |
| GRS1 | 1.20 | 9.99 |
| LSM4 | 1.58 | 9.79 |
| MFA2 | 1.80 | 9.62 |
| ACO1 | 1.12 | 9.57 |
| ARO9 | 1.23 | 9.53 |
| ACH1 | 1.56 | 9.48 |
| DGR1 | 2.45 | 9.45 |
| HHT2 | 1.01 | 9.44 |
| MCM3 | 1.28 | 9.41 |
| MAL12 | 2.14 | 9.40 |
| ERG1 | 1.09 | 9.36 |
| ADK1 | 1.22 | 9.29 |
| HIS4 | 1.23 | 9.27 |
| UTP8 | 1.30 | 9.25 |
| CIC1 | 1.03 | 9.13 |
| IRC13 | 1.40 | 9.12 |
| PWP1 | 1.25 | 9.10 |
| CGR1 | 1.29 | 9.01 |
| MDH1 | 1.06 | 9.01 |
| SKN1 | 1.28 | 8.98 |
| SDA1 | 1.08 | 8.91 |
| LSC2 | 1.03 | 8.91 |
| TAL1 | 1.19 | 8.89 |
| PCS60 | 1.07 | 8.87 |

| | | |
|---|---|---|
| PHO84 | 1.15 | 8.86 |
| AIM29 | 1.49 | 8.83 |
| SRP68 | 1.37 | 8.76 |
| TPM2 | 1.50 | 8.57 |
| GIR2 | 1.07 | 8.56 |
| MEU1 | 1.00 | 8.52 |
| COX4 | 1.06 | 8.50 |
| MHO1 | 1.15 | 8.35 |
| NOP12 | 1.00 | 8.35 |
| SMD3 | 1.38 | 8.31 |
| VAS1 | 1.04 | 8.20 |
| RLI1 | 1.21 | 8.15 |
| ISF1 | 1.12 | 8.10 |
| YGR067C | 1.59 | 8.05 |
| SNU13 | 1.06 | 8.02 |
| STL1 | 1.64 | 8.00 |
| SCP160 | 1.90 | 7.97 |
| MOG1 | 1.69 | 7.95 |
| HMO1 | 1.02 | 7.93 |
| YCL048W-A | 1.44 | 7.88 |
| YJL136W-A | 2.24 | 7.85 |
| BUD21 | 1.09 | 7.80 |
| YDR341C | 1.18 | 7.79 |
| WHI2 | 1.06 | 7.78 |
| LTP1 | 1.34 | 7.70 |
| PRP43 | 1.07 | 7.68 |
| YHR050W-A | 1.01 | 7.63 |
| TRX1 | 1.85 | 7.61 |
| LSM1 | 1.26 | 7.61 |
| ACS2 | 1.53 | 7.48 |
| ERG13 | 1.12 | 7.48 |
| DAP1 | 1.03 | 7.41 |
| CTI6 | 1.05 | 7.38 |
| CRF1 | 1.23 | 7.27 |
| TRM82 | 1.12 | 7.16 |
| BUR6 | 1.06 | 7.14 |
| YCR043C | 1.21 | 7.12 |

| | | |
|---|---|---|
| PPH3 | 1.24 | 6.97 |
| HOM3 | 1.33 | 6.96 |
| CDS1 | 1.25 | 6.88 |
| CBC2 | 1.23 | 6.87 |
| PCL7 | 1.09 | 6.85 |
| YKE2 | 1.47 | 6.75 |
| RSA1 | 1.28 | 6.67 |
| RVB1 | 1.01 | 6.67 |
| MCM5 | 1.50 | 6.66 |
| YOL038C-A | 2.43 | 6.63 |
| UBP13 | 1.11 | 6.62 |
| CLU1 | 1.22 | 6.60 |
| ELO3 | 1.40 | 6.56 |
| SPB1 | 1.02 | 6.52 |
| NCL1 | 1.12 | 6.41 |
| GPA1 | 1.32 | 6.40 |
| ADE2 | 1.12 | 6.40 |
| MIC10 | 1.01 | 6.39 |
| NDI1 | 1.04 | 6.34 |
| PUS4 | 1.88 | 6.30 |
| APS3 | 1.44 | 6.26 |
| YOR342C | 1.20 | 6.22 |
| UTP14 | 1.07 | 6.21 |
| FRD1 | 1.67 | 6.20 |
| RPA43 | 1.44 | 6.16 |
| RPC31 | 1.29 | 6.14 |
| DSD1 | 1.15 | 6.13 |
| GIM3 | 1.35 | 6.12 |
| MCM7 | 1.22 | 6.10 |
| CDC3 | 1.22 | 6.09 |
| YNCM0001W | 1.14 | 6.01 |
| SAP185 | 1.25 | 6.00 |
| MPP6 | 1.29 | 5.97 |
| CMS1 | 1.27 | 5.97 |
| DMO1 | 1.08 | 5.97 |
| PSD1 | 1.40 | 5.94 |
| ACL4 | 1.18 | 5.91 |

| | | |
|---|---|---|
| PRY1 | 1.93 | 5.85 |
| YCK2 | 1.12 | 5.82 |
| MIC60 | 1.37 | 5.74 |
| MKT1 | 1.07 | 5.74 |
| YER138W-A | 3.17 | 5.73 |
| VMA5 | 1.08 | 5.73 |
| PRO1 | 1.22 | 5.72 |
| NOP19 | 1.17 | 5.69 |
| PHM8 | 1.08 | 5.67 |
| FOL1 | 1.48 | 5.63 |
| END3 | 1.13 | 5.61 |
| USB1 | 1.79 | 5.60 |
| TSR1 | 1.20 | 5.55 |
| ARF3 | 1.45 | 5.52 |
| CKA1 | 1.00 | 5.46 |
| YNL162W-A | 1.08 | 5.45 |
| DAL2 | 1.67 | 5.43 |
| SCW4 | 1.74 | 5.39 |
| SNR39B | 1.27 | 5.34 |
| TCD2 | 1.10 | 5.32 |
| ALG5 | 1.65 | 5.28 |
| YGK1 | 1.10 | 5.25 |
| NUP57 | 1.09 | 5.25 |
| RPC11 | 1.15 | 5.21 |
| CYB2 | 1.72 | 5.20 |
| YEL068C | 1.62 | 5.17 |
| YIG1 | 1.15 | 5.15 |
| ARG4 | 1.03 | 5.11 |
| CIR2 | 1.12 | 5.06 |
| STE6 | 1.48 | 5.03 |
| BNA1 | 1.04 | 5.02 |
| PUS7 | 1.09 | 5.01 |
| RUD3 | 1.67 | 4.99 |
| RKM3 | 1.38 | 4.99 |
| YNL050C | 1.16 | 4.96 |
| YRO2 | 1.22 | 4.90 |
| TEA1 | 1.16 | 4.90 |

| | | |
|---|---|---|
| SME1 | 1.13 | 4.89 |
| TRM10 | 1.55 | 4.87 |
| RPC53 | 1.16 | 4.85 |
| YCR090C | 1.52 | 4.83 |
| REG2 | 1.01 | 4.82 |
| NOP9 | 1.24 | 4.73 |
| ACO2 | 1.01 | 4.69 |
| PLP2 | 1.08 | 4.67 |
| YNG1 | 1.11 | 4.61 |
| YCR016W | 1.22 | 4.58 |
| RRP1 | 1.17 | 4.58 |
| TSR3 | 1.01 | 4.53 |
| MAL32 | 1.68 | 4.50 |
| EMC10 | 1.03 | 4.49 |
| AIM7 | 1.56 | 4.39 |
| IPI3 | 1.05 | 4.37 |
| ENT5 | 1.08 | 4.33 |
| SEO1 | 1.74 | 4.31 |
| SHH4 | 1.14 | 4.31 |
| PER33 | 1.42 | 4.26 |
| DEG1 | 1.21 | 4.24 |
| CCW14 | 1.22 | 4.20 |
| SWC7 | 1.31 | 4.03 |
| YGR017W | 1.11 | 4.01 |
| HTB2 | 1.07 | 4.00 |
| SDO1 | 1.29 | 3.99 |
| CRS1 | 1.20 | 3.98 |
| HSP33 | 1.20 | 3.96 |
| SRP72 | 1.03 | 3.93 |
| ICL2 | 1.03 | 3.92 |
| SCC2 | 1.32 | 3.86 |
| RTT10 | 1.21 | 3.79 |
| PXP1 | 1.17 | 3.79 |
| BMT5 | 1.02 | 3.79 |
| NEJ1 | 1.45 | 3.78 |
| NMA1 | 1.20 | 3.76 |
| HSP32 | 1.20 | 3.74 |

| | | |
|---|---|---|
| NUP120 | 1.34 | 3.69 |
| DAS2 | 1.22 | 3.64 |
| ANT1 | 1.47 | 3.62 |
| AFR1 | 1.36 | 3.62 |
| JJJ3 | 1.13 | 3.62 |
| RRP45 | 1.06 | 3.60 |
| BET5 | 1.17 | 3.59 |
| GFD2 | 1.39 | 3.57 |
| MAK11 | 1.21 | 3.56 |
| DET1 | 1.05 | 3.56 |
| PXR1 | 1.07 | 3.45 |
| DUT1 | 1.27 | 3.44 |
| YJR056C | 1.20 | 3.44 |
| YNL035C | 1.11 | 3.44 |
| CSM4 | 1.20 | 3.43 |
| FMP23 | 1.15 | 3.43 |
| SGN1 | 1.15 | 3.43 |
| GIP1 | 1.43 | 3.42 |
| UPF3 | 1.17 | 3.42 |
| HGH1 | 1.10 | 3.40 |
| VPS63 | 1.29 | 3.35 |
| YVH1 | 1.03 | 3.34 |
| FAL1 | 1.40 | 3.33 |
| SAM4 | 1.25 | 3.33 |
| MDE1 | 1.13 | 3.33 |
| FKH1 | 1.39 | 3.30 |
| NAT5 | 1.05 | 3.24 |
| JJJ1 | 1.06 | 3.21 |
| YOS9 | 1.14 | 3.14 |
| NRP1 | 1.11 | 3.13 |
| DAD3 | 1.15 | 3.12 |
| PRO3 | 1.09 | 3.12 |
| RPC25 | 1.03 | 3.10 |
| SEC12 | 1.02 | 3.08 |
| SAP30 | 1.11 | 3.06 |
| YNCP0019W | 1.35 | 3.03 |
| LDO45 | 1.04 | 3.03 |

| | | |
|---|---|---|
| EPT1 | 1.68 | 3.00 |
| MNT2 | 1.34 | 2.98 |
| MLP2 | 1.03 | 2.97 |
| NRK1 | 1.46 | 2.96 |
| GPI15 | 1.14 | 2.95 |
| FMP41 | 1.03 | 2.93 |
| INM1 | 1.05 | 2.92 |
| RFC1 | 1.26 | 2.91 |
| NAT1 | 1.04 | 2.90 |
| FMP32 | 1.01 | 2.88 |
| YDL086W | 1.05 | 2.83 |
| SPO12 | 1.49 | 2.67 |
| TAF11 | 1.10 | 2.66 |
| YEL067C | 1.16 | 2.60 |
| SWI6 | 1.30 | 2.57 |
| PIR1 | 1.03 | 2.55 |
| YGL081W | 1.71 | 2.53 |
| TRM3 | 1.06 | 2.53 |
| YNCG0008W | 2.49 | 2.52 |
| SOL3 | 1.21 | 2.52 |
| BMT6 | 1.37 | 2.51 |
| CAF40 | 1.07 | 2.46 |
| BBP1 | 1.65 | 2.42 |
| SEN34 | 1.52 | 2.36 |
| DCD1 | 1.82 | 2.34 |
| RMP1 | 1.25 | 2.33 |
| YOL029C | 1.01 | 2.26 |
| YGR240C-A | 1.44 | 2.21 |
| LYS9 | 1.06 | 2.19 |
| IOC3 | 1.00 | 2.19 |
| ASH1 | 1.72 | 2.14 |
| ECM9 | 1.25 | 2.13 |
| NKP1 | 1.30 | 2.11 |
| YAP7 | 1.05 | 2.07 |
| ADH4 | 1.01 | 2.06 |
| GID10 | 1.20 | 2.02 |
| PIG1 | 1.24 | 2.01 |

| | | |
|---|---|---|
| YBR238C | 1.13 | 2.01 |
| MDM12 | 1.02 | 2.01 |

[a]UniProt (https://www.uniprot.org/)

[b]*Saccharomyces* Genome Database (https://www.yeastgenome.org/)

**Supplementary Table 4 Genes in yeast downregulated via co-cultivation**

| Gene[a] or locus tag[b] | $Log_2$ (fold change) | $-Log_{10}$ (p-value) |
|---|---|---|
| YPR064W | -4.12 | 3.26 |
| SAC6 | -1.02 | 18.84 |
| YPR145C-A | -1.50 | 18.17 |
| YDL012C | -1.60 | 17.98 |
| BAP3 | -1.51 | 17.92 |
| YLR108C | -2.21 | 17.66 |
| RSN1 | -1.29 | 17.47 |
| ARK1 | -1.58 | 17.43 |
| ASE1 | -1.42 | 17.43 |
| TVP18 | -1.56 | 17.41 |
| YAR029W | -2.38 | 17.32 |
| PRC1 | -1.15 | 16.96 |
| FET5 | -1.54 | 16.76 |
| YDR210W | -1.95 | 16.60 |
| NUM1 | -1.56 | 16.60 |
| YHR214C-C | -1.41 | 16.49 |
| PIL1 | -1.35 | 16.31 |
| FLC2 | -1.08 | 16.31 |
| SNR81 | -1.96 | 16.28 |
| YML040W | -1.41 | 16.25 |
| YHR139C-A | -2.68 | 16.19 |
| OYE3 | -1.68 | 16.02 |
| YOL103W-A | -1.41 | 15.81 |
| GAT2 | -2.19 | 15.78 |
| THP3 | -1.51 | 15.75 |
| FMP45 | -1.24 | 15.63 |
| tP(UGG)Q | -3.06 | 15.60 |
| HUR1 | -1.43 | 15.41 |
| SNR70 | -1.01 | 15.22 |
| SSD1 | -1.43 | 15.20 |
| PMU1 | -1.86 | 15.03 |
| YLL053C | -3.28 | 14.93 |
| MDJ1 | -1.24 | 14.89 |
| YHK8 | -1.98 | 14.76 |
| YSR3 | -1.86 | 14.74 |

| | | |
|---|---|---|
| YLR227W-A | -1.34 | 14.69 |
| YGL088W | -1.01 | 14.69 |
| YOL036W | -1.75 | 14.57 |
| GID8 | -1.13 | 14.52 |
| CCP1 | -1.42 | 14.49 |
| YHL015W-A | -2.66 | 14.39 |
| YNL234W | -1.76 | 14.39 |
| GSH1 | -1.13 | 14.34 |
| YBR016W | -1.57 | 14.32 |
| LIH1 | -1.84 | 14.20 |
| AQY2 | -3.21 | 14.04 |
| MEO1 | -1.06 | 14.01 |
| MFG1 | -1.94 | 13.95 |
| YPR158C-C | -1.35 | 13.93 |
| MIM2 | -1.86 | 13.92 |
| RFS1 | -1.39 | 13.85 |
| PDR15 | -1.82 | 13.54 |
| PEP12 | -1.05 | 13.50 |
| APJ1 | -1.32 | 13.40 |
| CPR6 | -1.06 | 13.36 |
| TOM6 | -1.30 | 13.28 |
| THO2 | -1.72 | 13.22 |
| RDN5-1 | -3.48 | 13.17 |
| ITS2-2 | -1.19 | 13.11 |
| SAF1 | -1.57 | 12.96 |
| ECM27 | -2.03 | 12.89 |
| PMR1 | -1.07 | 12.88 |
| AGP2 | -1.67 | 12.77 |
| TOG1 | -1.55 | 12.77 |
| PHM7 | -1.52 | 12.66 |
| ZRT3 | -1.61 | 12.64 |
| LTO1 | -1.95 | 12.48 |
| YDR215C | -2.46 | 12.29 |
| CSR2 | -1.13 | 12.27 |
| YGR050C | -2.24 | 12.17 |
| MCA1 | -1.34 | 12.15 |
| LDS2 | -2.25 | 12.06 |

| | | |
|---|---|---|
| YPR158W-A | -1.47 | 12.04 |
| YNL054W-A | -1.30 | 11.71 |
| PSK2 | -1.36 | 11.67 |
| RCN1 | -1.85 | 11.60 |
| LGE1 | -1.72 | 11.59 |
| YLR162W | -1.50 | 11.48 |
| COS6 | -1.04 | 11.48 |
| YGR174W-A | -1.46 | 11.44 |
| SIS1 | -1.02 | 11.33 |
| NBP35 | -1.18 | 11.29 |
| YMR315W-A | -2.33 | 11.25 |
| UGA2 | -1.01 | 11.24 |
| MDY2 | -1.55 | 11.15 |
| YFL051C | -1.44 | 11.11 |
| YBR284W | -2.20 | 11.10 |
| HBN1 | -2.30 | 10.95 |
| LYS20 | -1.41 | 10.91 |
| VMA11 | -1.49 | 10.88 |
| RIM8 | -1.04 | 10.87 |
| SHE10 | -1.47 | 10.83 |
| TOS8 | -1.11 | 10.82 |
| YPL278C | -2.42 | 10.79 |
| ENB1 | -1.56 | 10.74 |
| CSS1 | -1.53 | 10.74 |
| HXT10 | -2.40 | 10.53 |
| SPF1 | -1.14 | 10.52 |
| RPL29 | -1.02 | 10.45 |
| ZWF1 | -1.44 | 10.43 |
| MIN3 | -1.18 | 10.41 |
| YGR038C-A | -1.39 | 10.37 |
| MNN14 | -1.34 | 10.37 |
| RCH1 | -1.22 | 10.33 |
| ATG1 | -1.26 | 10.31 |
| YNCK0021W | -3.65 | 10.28 |
| UBP5 | -1.42 | 10.27 |
| MSP1 | -1.16 | 10.25 |
| YCR102C | -2.38 | 10.24 |

| | | |
|---|---|---|
| SNR49 | -1.16 | 10.15 |
| HXT8 | -2.57 | 10.13 |
| YNL018C | -2.65 | 10.11 |
| AUA1 | -1.18 | 10.07 |
| TPH3 | -1.52 | 10.04 |
| VRG4 | -1.15 | 10.01 |
| PRM5 | -1.23 | 9.95 |
| YLR345W | -1.39 | 9.92 |
| YOL118C | -1.39 | 9.92 |
| OPT1 | -1.63 | 9.90 |
| SNG1 | -1.48 | 9.74 |
| TPO2 | -2.10 | 9.71 |
| UGA4 | -1.09 | 9.68 |
| IRC20 | -1.01 | 9.59 |
| POM33 | -2.85 | 9.57 |
| DOA4 | -1.76 | 9.52 |
| LAP3 | -1.63 | 9.41 |
| YHR140W | -2.17 | 9.36 |
| YDL144C | -2.43 | 9.29 |
| YNL143C | -1.76 | 9.24 |
| YGR079W | -1.26 | 9.20 |
| FRE6 | -1.42 | 9.15 |
| AVT6 | -1.99 | 9.12 |
| YMR027W | -1.42 | 9.10 |
| YDC1 | -1.31 | 8.90 |
| RPL15B | -1.27 | 8.90 |
| IMA4 | -2.03 | 8.80 |
| YMR008C-A | -1.94 | 8.79 |
| PKH2 | -1.65 | 8.75 |
| SNR128 | -1.21 | 8.72 |
| SCT1 | -1.35 | 8.71 |
| YPL277C | -2.82 | 8.65 |
| SNR47 | -2.20 | 8.51 |
| TDA9 | -1.00 | 8.50 |
| RDR1 | -2.00 | 8.49 |
| YDR316W-A | -1.17 | 8.49 |
| UIP3 | -1.30 | 8.46 |

| | | |
|---|---|---|
| IMA3 | -2.12 | 8.45 |
| PET9 | -1.26 | 8.44 |
| VHS2 | -1.86 | 8.38 |
| YML045W-A | -1.20 | 8.33 |
| FLO11 | -2.21 | 8.19 |
| YGR161C-C | -1.15 | 8.19 |
| TYC1 | -1.53 | 8.14 |
| YDR098C-A | -1.20 | 8.10 |
| PCT1 | -1.12 | 8.07 |
| JSN1 | -1.66 | 8.05 |
| MHF2 | -1.12 | 8.05 |
| SKG3 | -1.71 | 7.94 |
| FET3 | -1.12 | 7.94 |
| ITS1-2 | -1.48 | 7.93 |
| YAL064W | -2.00 | 7.89 |
| PUL4 | -1.50 | 7.86 |
| CPR4 | -1.34 | 7.84 |
| CAD1 | -1.08 | 7.82 |
| KIN1 | -1.09 | 7.79 |
| CYT2 | -1.08 | 7.77 |
| YOR142W-A | -1.27 | 7.76 |
| ZNF1 | -1.18 | 7.69 |
| OSW7 | -1.09 | 7.64 |
| APP1 | -1.44 | 7.51 |
| YTP1 | -1.41 | 7.50 |
| MCO8 | -1.10 | 7.48 |
| PSK1 | -1.22 | 7.41 |
| FPK1 | -1.13 | 7.41 |
| SOM1 | -1.39 | 7.40 |
| YDR210C-C | -1.23 | 7.39 |
| YER159C-A | -1.16 | 7.30 |
| MSN4 | -1.32 | 7.28 |
| XBP1 | -1.87 | 7.27 |
| IRC10 | -1.99 | 7.25 |
| MRPL35 | -1.21 | 7.20 |
| SNR48 | -2.90 | 7.19 |
| HPF1 | -1.02 | 7.19 |

| | | |
|---|---|---|
| YOR192C-A | -1.74 | 7.16 |
| MGR3 | -1.56 | 7.16 |
| RRT1 | -1.22 | 7.10 |
| YBL100W-C | -2.10 | 7.09 |
| ATX1 | -1.33 | 7.00 |
| SMP1 | -1.12 | 6.95 |
| YOL163W | -1.61 | 6.90 |
| MNT4 | -1.17 | 6.84 |
| ATG9 | -1.17 | 6.84 |
| PTK1 | -2.52 | 6.83 |
| YMR051C | -1.09 | 6.83 |
| TGL4 | -1.31 | 6.81 |
| YNCL0033C | -3.60 | 6.80 |
| MRX16 | -1.28 | 6.80 |
| YAP5 | -1.00 | 6.79 |
| ECM19 | -2.84 | 6.78 |
| UBA4 | -1.19 | 6.70 |
| PAP2 | -1.48 | 6.56 |
| MER1 | -1.09 | 6.54 |
| YNL019C | -1.54 | 6.50 |
| LCL1 | -1.34 | 6.48 |
| AKL1 | -1.42 | 6.46 |
| SHR5 | -1.07 | 6.46 |
| SUF2 | -1.96 | 6.39 |
| PTP2 | -1.24 | 6.36 |
| YLR415C | -1.91 | 6.30 |
| RRT6 | -1.24 | 6.30 |
| YNCG0011W | -3.03 | 6.26 |
| NGL3 | -1.18 | 6.26 |
| STE18 | -1.15 | 6.26 |
| ITS1-1 | -1.27 | 6.23 |
| YFT2 | -1.11 | 6.22 |
| MUP3 | -1.48 | 6.21 |
| ASI2 | -1.43 | 6.15 |
| SPC1 | -1.00 | 6.15 |
| IMA2 | -1.86 | 6.12 |
| YET3 | -1.01 | 6.08 |

| | | |
|---|---|---|
| TUL1 | -1.61 | 6.06 |
| SOK2 | -1.23 | 6.05 |
| VPS36 | -1.18 | 5.96 |
| YGR237C | -1.98 | 5.82 |
| GPI13 | -1.54 | 5.81 |
| MLO1 | -1.21 | 5.81 |
| MSC3 | -1.10 | 5.79 |
| ATG41 | -1.09 | 5.75 |
| YHL026C | -1.23 | 5.74 |
| ATR1 | -1.02 | 5.72 |
| YGL007C-A | -1.16 | 5.65 |
| STB3 | -1.55 | 5.63 |
| LYS14 | -1.36 | 5.62 |
| YBR012W-A | -1.14 | 5.61 |
| CDC16 | -1.60 | 5.59 |
| YNCO0021C | -2.89 | 5.48 |
| VPS71 | -1.33 | 5.47 |
| STP1 | -1.07 | 5.44 |
| NRD1 | -1.34 | 5.39 |
| SDS24 | -2.41 | 5.37 |
| OSW2 | -1.71 | 5.37 |
| ATG26 | -1.11 | 5.33 |
| YMR158C-A | -1.65 | 5.32 |
| SNX4 | -1.14 | 5.31 |
| YLR001C | -1.00 | 5.29 |
| YNCG0002C | -1.53 | 5.28 |
| AUR1 | -1.16 | 5.27 |
| ATG40 | -1.34 | 5.23 |
| AFT1 | -1.91 | 5.18 |
| YAT1 | -1.68 | 5.16 |
| AGE2 | -1.47 | 5.14 |
| MOR1 | -1.33 | 5.13 |
| GIS1 | -1.04 | 5.12 |
| RRG9 | -1.11 | 5.09 |
| YOL014W | -1.45 | 5.08 |
| SFK1 | -1.50 | 5.01 |
| FRE7 | -1.26 | 5.01 |

| | | |
|---|---|---|
| TIM13 | -1.36 | 5.00 |
| YFL002W-B | -1.44 | 4.95 |
| YIL055C | -1.72 | 4.91 |
| YSF3 | -1.26 | 4.87 |
| PPM1 | -1.17 | 4.84 |
| ROF1 | -2.24 | 4.83 |
| YGR161W-A | -1.35 | 4.82 |
| YMR141C | -1.06 | 4.79 |
| MRX14 | -1.02 | 4.79 |
| YNL140C | -1.37 | 4.78 |
| HMS2 | -1.03 | 4.77 |
| ICR1 | -1.79 | 4.76 |
| PUS2 | -1.93 | 4.71 |
| SAM35 | -1.60 | 4.69 |
| ASG7 | -1.48 | 4.67 |
| YOL131W | -2.37 | 4.64 |
| YOL162W | -1.55 | 4.62 |
| IMA1 | -2.29 | 4.60 |
| PAN1 | -1.72 | 4.59 |
| ATG2 | -1.09 | 4.56 |
| DUG3 | -1.25 | 4.50 |
| YJL077W-A | -1.01 | 4.49 |
| YML054C-A | -2.41 | 4.47 |
| MHP1 | -1.12 | 4.46 |
| MUM3 | -1.27 | 4.45 |
| YER158C | -1.26 | 4.44 |
| YJL163C | -1.39 | 4.41 |
| ECM21 | -1.12 | 4.41 |
| YNL284C-A | -1.30 | 4.40 |
| CUE4 | -1.04 | 4.39 |
| CDC27 | -1.20 | 4.33 |
| ALY2 | -1.58 | 4.32 |
| YBL086C | -1.54 | 4.31 |
| MLF3 | -1.38 | 4.30 |
| YOR365C | -1.47 | 4.28 |
| RFA3 | -1.46 | 4.24 |
| YIR020C | -1.74 | 4.23 |

| | | |
|---|---|---|
| YOR292C | -1.58 | 4.23 |
| KEX2 | -1.08 | 4.15 |
| YJL052C-A | -3.00 | 4.13 |
| HAP1 | -1.14 | 4.12 |
| INA22 | -1.01 | 4.11 |
| YGL193C | -1.49 | 4.09 |
| TOP3 | -2.09 | 4.07 |
| AHC1 | -1.49 | 4.07 |
| tE(UUC)Q | -2.04 | 4.01 |
| PSF3 | -1.25 | 3.99 |
| CHL4 | -1.10 | 3.99 |
| BOL3 | -1.41 | 3.98 |
| CAB3 | -1.01 | 3.97 |
| JAC1 | -1.52 | 3.96 |
| YIL163C | -2.33 | 3.94 |
| FIP1 | -1.51 | 3.94 |
| YIL102C-A | -1.03 | 3.94 |
| IXR1 | -1.35 | 3.92 |
| UPS3 | -1.19 | 3.91 |
| TMS1 | -1.16 | 3.91 |
| SED5 | -1.49 | 3.89 |
| PAU4 | -3.33 | 3.88 |
| KEL1 | -1.35 | 3.88 |
| LAT1 | -1.32 | 3.88 |
| BBC1 | -1.42 | 3.86 |
| RTA1 | -1.46 | 3.80 |
| ASK1 | -1.02 | 3.77 |
| PRM1 | -1.49 | 3.73 |
| YLR125W | -1.23 | 3.73 |
| YNCE0009C | -2.59 | 3.72 |
| VFA1 | -1.77 | 3.71 |
| NBA1 | -1.50 | 3.71 |
| SNR52 | -1.23 | 3.71 |
| YLR177W | -1.07 | 3.68 |
| ACK1 | -1.42 | 3.67 |
| HER1 | -1.28 | 3.67 |
| YML100W-A | -1.88 | 3.66 |

| | | |
|---|---|---|
| MGA1 | -1.58 | 3.63 |
| YLR236C | -1.88 | 3.62 |
| PWR1 | -2.16 | 3.58 |
| NVJ2 | -1.14 | 3.57 |
| YOR389W | -1.31 | 3.56 |
| ACE2 | -1.76 | 3.55 |
| CDC13 | -1.54 | 3.52 |
| MYO3 | -1.16 | 3.52 |
| YGR266W | -1.27 | 3.49 |
| YRR1 | -1.03 | 3.48 |
| GTO3 | -1.02 | 3.48 |
| MCT1 | -1.02 | 3.48 |
| BUD7 | -1.02 | 3.47 |
| MRS1 | -1.54 | 3.41 |
| MET13 | -1.13 | 3.41 |
| YLR046C | -1.19 | 3.39 |
| YDR102C | -1.72 | 3.38 |
| YSC84 | -1.36 | 3.38 |
| FRE8 | -1.12 | 3.37 |
| PEX6 | -1.02 | 3.37 |
| YNCP0010W | -2.46 | 3.36 |
| QDR1 | -1.11 | 3.36 |
| IME4 | -1.56 | 3.32 |
| KAR3 | -1.18 | 3.30 |
| YLR412C-A | -2.23 | 3.29 |
| DSC2 | -1.25 | 3.28 |
| YHR214C-D | -2.62 | 3.26 |
| BSP1 | -1.65 | 3.24 |
| YNCG0028W | -2.63 | 3.23 |
| YPR084W | -1.09 | 3.23 |
| GPT2 | -1.08 | 3.21 |
| RME2 | -1.54 | 3.20 |
| GEP7 | -1.41 | 3.18 |
| VID28 | -1.08 | 3.18 |
| GAL80 | -1.19 | 3.14 |
| KTR3 | -1.13 | 3.07 |
| YOR343C | -1.68 | 3.06 |

| | | |
|---|---|---|
| SNR6 | -2.75 | 3.05 |
| TRK2 | -1.01 | 3.04 |
| tF(GAA)Q | -1.47 | 3.03 |
| MIX14 | -1.03 | 3.03 |
| MOT2 | -1.04 | 3.02 |
| APM2 | -1.67 | 3.00 |
| YDL109C | -1.48 | 2.99 |
| YOR032W-A | -1.11 | 2.99 |
| YDR114C | -1.54 | 2.97 |
| YIL102C | -1.86 | 2.96 |
| PAU23 | -2.16 | 2.92 |
| EAP1 | -2.06 | 2.92 |
| YOR387C | -1.86 | 2.91 |
| YNL033W | -1.33 | 2.90 |
| SLZ1 | -1.55 | 2.87 |
| SHC1 | -1.35 | 2.87 |
| YJL043W | -2.20 | 2.84 |
| SNR58 | -1.46 | 2.84 |
| DIN7 | -1.12 | 2.84 |
| RTT105 | -1.11 | 2.83 |
| YNL277W-A | -2.19 | 2.80 |
| PHR1 | -1.56 | 2.78 |
| THI74 | -1.05 | 2.76 |
| YNCP0009W | -1.52 | 2.73 |
| YKR073C | -1.32 | 2.73 |
| MGL2 | -1.08 | 2.73 |
| PAM17 | -1.07 | 2.73 |
| YJL213W | -1.69 | 2.72 |
| YLL056C | -1.09 | 2.68 |
| YIL060W | -1.13 | 2.66 |
| BET2 | -1.00 | 2.66 |
| DIE2 | -1.73 | 2.65 |
| CIN10 | -1.16 | 2.65 |
| ADY3 | -1.19 | 2.64 |
| MSN2 | -1.03 | 2.63 |
| YIL171W | -1.45 | 2.62 |
| RMR1 | -1.20 | 2.61 |

| | | |
|---|---|---|
| GEM1 | -1.07 | 2.61 |
| SEC16 | -1.35 | 2.60 |
| IRC22 | -1.04 | 2.59 |
| MMS4 | -1.08 | 2.58 |
| RNR3 | -1.33 | 2.57 |
| YNCM0029C | -2.27 | 2.55 |
| MED2 | -1.08 | 2.55 |
| YAR035C-A | -1.30 | 2.53 |
| ROG3 | -1.09 | 2.53 |
| YHR130C | -1.04 | 2.52 |
| SNR62 | -1.02 | 2.50 |
| NIS1 | -1.01 | 2.50 |
| SLO1 | -1.03 | 2.48 |
| YNCG0004W | -2.46 | 2.44 |
| YNCC0005W | -1.83 | 2.43 |
| OST3 | -1.53 | 2.35 |
| VLD1 | -1.07 | 2.32 |
| VHR1 | -1.06 | 2.32 |
| YMR272W-B | -2.41 | 2.28 |
| PGU1 | -1.71 | 2.27 |
| SNR64 | -1.31 | 2.27 |
| COA4 | -1.01 | 2.27 |
| YGR109W-A | -1.63 | 2.25 |
| RPI1 | -1.35 | 2.24 |
| CIN5 | -1.09 | 2.23 |
| AMF1 | -1.03 | 2.20 |
| YHC3 | -1.54 | 2.19 |
| NTE1 | -1.19 | 2.18 |
| AIM3 | -1.26 | 2.14 |
| WSC2 | -1.14 | 2.14 |
| YPL080C | -1.14 | 2.14 |
| PFS2 | -1.03 | 2.14 |
| YDL186W | -3.62 | 2.10 |
| FRE5 | -1.51 | 2.09 |
| MRX11 | -1.35 | 2.09 |
| YKR045C | -1.10 | 2.09 |
| YLR406C-A | -2.68 | 2.08 |

| | | |
|---|---|---|
| YOR268C | -2.13 | 2.06 |
| SPR2 | -1.07 | 2.06 |
| ZRG8 | -1.47 | 2.02 |

[a]UniProt (https://www.uniprot.org/)

[b]*Saccharomyces* Genome Database (https://www.yeastgenome.org/)
